# Identifying Regulation with Adversarial Surrogates

**DOI:** 10.1101/2022.10.08.511451

**Authors:** Ron Teichner, Aseel Shomar, O. Barak, N. Brenner, S. Marom, R. Meir, D. Eytan

**Author notes:** Equal contribution (Ron Teichner, Aseel Shomar).

## Abstract

Homeostasis, the ability to maintain a relatively constant internal environment in the face of perturbations, is a hallmark of biological systems. It is believed that this constancy is achieved through multiple internal regulation and control processes. Given observations of a system, or even a detailed model of one, it is both valuable and extremely challenging to extract the control objectives of the homeostatic mechanisms. In this work, we develop a robust data-driven method to identify these objectives, namely to understand: “what does the system care about?”. We propose an algorithm, Identifying Regulation with Adversarial Surrogates (IRAS), that receives an array of temporal measurements of the system, and outputs a candidate for the control objective, expressed as a combination of observed variables. IRAS is an iterative algorithm consisting of two competing players. The first player, realized by an artificial deep neural network, aims to minimize a measure of invariance we refer to as the coefficient of regulation. The second player aims to render the task of the first player more difficult by forcing it to extract information about the temporal structure of the data, which is absent from similar ‘surrogate’ data. We test the algorithm on two synthetic and one natural data set, demonstrating excellent empirical results. Interestingly, our approach can also be used to extract conserved quantities, e.g., energy and momentum, in purely physical systems, as we demonstrate empirically.

## 1 Introduction

Living systems maintain stability against internal and external perturbations, a phenomenon known as homeostasis [Billman, 2020, Hsiao et al., 2018, Kotas and Medzhitov, 2015]. This is a ubiquitous central pillar across all scales of biological organization, such as molecular circuits, physiological functions, and population dynamics. Failure of homeostatic control is associated with diseases including diabetes, autoimmunity, and obesity [Kotas and Medzhitov, 2015]. It is therefore vital to identify the regulated variables that the system aims to maintain at a stable setpoint.

In contrast to simple human-made systems, where often a small number of known variables are under control, biological systems are characterized by multiple coupled feedback loops as well as other dynamic structures [Billman, 2020]. In particular, they are not divided to separate ‘plant’ and ‘controller’ entities, as commonly characterized in control theory, but rather make up a complex network of interactions. In such a network some variables are maintained at a stable setpoint, whereas others are more flexibly modulated to maintain the former regulated variables around their setpoints. A classic example is blood glucose concentration, which is tightly regulated, while the rates of glycolysis and gluconeogenesis are flexible variables [Kotas and Medzhitov, 2015]. Thus, in general one may find a hierarchy of control, where some variables are more tightly controlled than others. [El-Samad, 2021, Stawsky et al., 2021]. This biological complexity makes it challenging to identify the regulated variables that the system actively maintains at a setpoint, in contrast to those which are stabilized as a byproduct.

Experimentally, a regulated variable can be identified by performing perturbations [Tegnér and Björkegren, 2007]. When the system is perturbed, compensatory mechanisms will adjust other variables to restore it to its setpoint by using feedback [El-Samad, 2021]. However, designing such experiments requires prior knowledge about the system, which is not always at hand and may be technically challenging or infeasible. Biological systems regulate internal variables, rather than measured variables, which are generally determined by experimental constraints. Our assumption, related to the concept of observability [Liu et al., 2013], is that a *combination* of the observed variables will correspond to the relevant internal variable.

In recent years, technological advancements brought about a huge number of available datasets that were not tailored to find regulated variables, but could offer the opportunity to point out possible candidates. This raises the question: can we elicit the regulated variables of a system given a set of measurements with minimal prior assumptions?

In this work, we develop an algorithm, Identifying Regulation with Adversrial Surrogates (IRAS), that aims to identify the most conserved combination of variables in a system. This combination, operationally denoted the control objective, may represent a quantity that is of high importance to the system. To this end, a quantitative measure needs to be defined, which enables comparing the degree of invariance of different combinations. Standard statistical measures, such as the variance or coefficient of variation, are not suitable due to their sensitivity to scale and bias, and insensitivity to temporal aspects. We propose a new measure, the Coefficient of Regulation (CR), which captures the property of temporal invariance. Straightforward optimization of this measure does not provide the required result (for reasons explained below). Rather, we show that a combined utilization of temporal invariance, and the geometric distribution of data, can be successful in the task.

IRAS receives as input an array of temporal measurements, and outputs the control objective as a combination (function) of the observed variables. At its core, it runs iteratively between two competing players; one player aims to minimize the CR, i.e. to find the combination which is most invariant in the data relative to time-shuffled data. The second player gradually pushes the time-shuffled ensemble to statistically resemble the real data, thus rendering the optimization problem more difficult for the first player. Eventually, the control objective is found when these two players converge.

To demonstrate the generality of our approach, we validate it on three examples from very different domains. First, we simulate a kinetic model of protein-protein interactions, in which the controlled variable combination is known. We show that IRAS identifies with high accuracy the control objective. Second, we analyze data from a psychophysical experiment, where human observers’ response statistics are modulated by an artificial controller. Our algorithm identifies correctly the known control circuit. Finally, we leave the biological domain and illustrate the generality of IRAS by considering an example of energy conservation in a physical spring system. We demonstrate that, based on observing noisy trajectories of multiple spring systems with different parameters, we are able to recover both the individual parameters of each system and the ‘law of conservation of energy’, with an explicit form for the energy function. To the best of our knowledge, such an empirical approach has not been developed previously, and could potentially be useful to many experimental systems.

### Problem illustration

We are interested in identifying empirically, from a set of measurements, a quantity which is most conserved around a setpoint. This “control objective” could represent something of high importance to the system and could thus shed light on the system’s functionality. How can we elicit a possible control objective of a system given a set of *n* measurements over time, *Z*(*t*) = (*z*_1_(*t*), …, *z*_*n*_(*t*))? If the control objective itself is unknown, it is likely not directly measured. However, it could be possible to describe it as a combination of the measured variables, that is maintained around a setpoint,

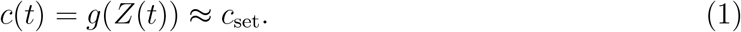

As a simple illustrative example, consider the case of three proteins whose abundance is measured across time in a single cell. Figure 1A shows these traces along time, as the system presumably goes through various perturbations. While the amount of the three proteins, *P*_1_(*t*), *P*_2_(*t*) and *P*_3_(*t*), varies significantly over time, the ratio 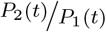 is maintained around a setpoint *c*_set_ = 2 with small fluctuations (black line). No other instantaneous relationship between the proteins is present in the data. We thus expect the combination 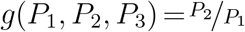 to be identified as the control objective of the system.

**Figure 1:**
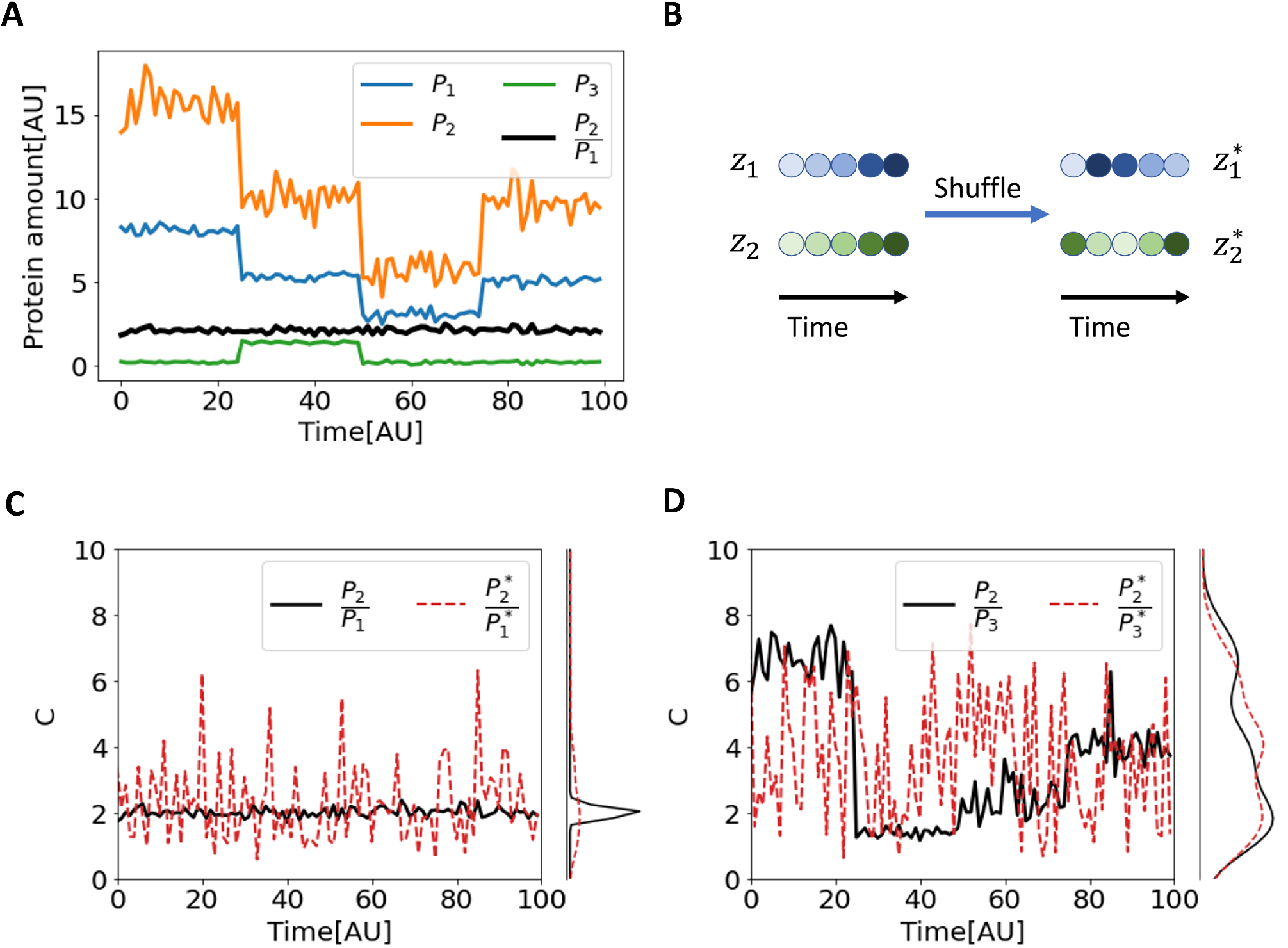
Coefficient of Regulation (CR) as a measure of the combination invariance. **(A)** A synthetic example of ratio control in a biological system. The amounts of three proteins, *P*_1_, *P*_2_ and *P*_3_, fluctuate over time and are modulated by discontinuous perturbations (*t* = 25, 50, 75). In the face of these perturbations, the ratio 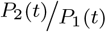 is held around a setpoint *c*_*set*_ = 2 with small fluctuations (noise with standard deviation 0.15). **(B)** To calculate CR, temporal correlations between the measurements are destroyed by shuffling the time points of each measurement independently. **(C)** While the distribution of 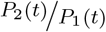 in real data is narrow (black), the distribution of the ratio between the shuffled measurements is much wider (red dashed), resulting in a small CR. **(D)** Since there is no correlation between *P*_2_ and *P*_3_, the distributions of their ratio in real and in shuffled data are of the same width, and CR= 1.

Our aim in what follows is to develop an algorithm that receives as input a set of experimental measurements, and outputs the control objective as a combination of current, and possibly previous, measurements. After defining a measure of invariance (Section 2.1), we explain why simply optimizing it is insufficient (2.2) and construct IRAS, a two-player algorithm, to minimize it under iterative constraints (2.3). Then, we validate the algorithm on three examples, where the control objective depends on current variables in one case (3.1), extended to include also past values in a second case (3.2) and then also extended to include system-specific parameter estimation in the third case (3.3). IRAS is data driven, and is not provided with a model of the system, or with possible candidates for the control objective based on prior knowledge. Rather, it is based solely on the raw measurements.

## 2 Algorithm Development

### 2.1 Quantifying invariance around a setpoint

Based on our assumption that a regulated variable is held relatively constant, we first seek a measure that quantifies the invariance of a combination around a stable setpoint. We posit that the controller couples system variables (such as the levels of the two proteins above) that would otherwise be less, or even completely, decoupled. As perturbations are encountered, these variables co-vary, and their joint distribution

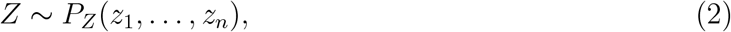

defines the geometry of the manifold which the data occupy. By arbitrarily permuting each component *z*_*i*_ independently over time, we can create a surrogate data-set *Z*^*^ in which the correlations in the data have been eliminated – in particular those induced by the control. The distribution of this surrogate data-set reflects only the single-variable properties:

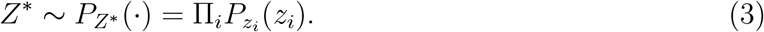

Importantly, a combination *c*(*t*) = *g* (*Z*(*t*)) that is invariant due to the operation of the controller, would become non-invariant in the surrogate data,

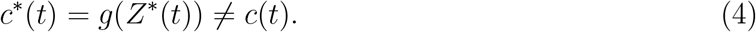

To quantify the sensitivity of a combination to independent shuffling, we consider the ratio

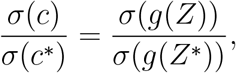

where *σ* is the standard deviation computed over time. We expect that for a regulated combination, destroying all temporal order will increase its standard deviation considerably, and decrease this ratio. Note that when the components of *Z* are independent, i.e. there is no relation between them, the ratio is identically 1 for any combination *g*.

Figure 1B illustrates this definition: starting from measurements *Z*, we create the independently shuffled ensemble *Z*^*^ where correlations between variables are destroyed. Referring to the protein example in Figure 1A, the combination 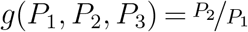 is maintained around the setpoint, resulting in a narrow distribution over time and a small standard deviation (Figure 1C, black). Eliminating the temporal correlation between *P*_1_ and *P*_2_ by shuffling their time points, results in a much wider distribution and a higher standard deviation (Figure 1C, dashed red).Other combinations such as 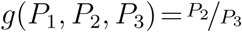, exhibit distributions of similar widths over the shuffled and the original data (Figure 1D). Consequently, the standard deviation ratio is approximately 1, indicating that this combination is not regulated by the system.

The standard deviation ratio, that quantifies invariance in temporally-ordered vs. temporally-shuffled data, can be generalized to shuffles that are not completely random but obey some constraint. As shown below, this generalization will be required for developing our two-player algorithm. Specifically, given a suggested combination *c* = *g*(·), one may construct a weighting function *ζ*(·), that defines a biased shuffled ensemble 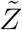. Then, we define the Coefficient of Regulation (CR) as follows:

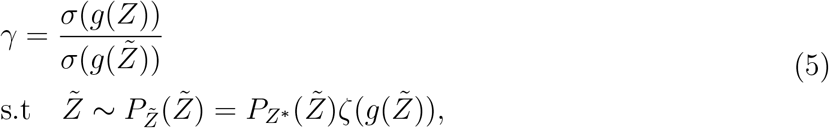

where the special case of completely random shuffles is obtained for *ζ* = 1.

To summarize, we defined the Coefficient of Regulation (CR) as a measure that quantifies the sensitivity of a given combination to the destruction of temporal correlations between its constituents. It is based on a surrogate data technique [Lancaster et al., 2018], applied here to multiple variables by shuffling each of them separately, and measuring the effect on their combinations. The shuffles can be completely random or performed under some constraints. We next consider the question of how this measure can be used to identify the control objective without prior assumptions.

### 2.2 Straightforward optimization fails by shuffle artefacts

Since low values of CR indicate invariance around a setpoint, one may expect that the combination that minimizes CR with respect to unconstrained shuffling, *ζ*(·) ≡ 1 in (5), is a good candidate for the control objective of the system. If so, we would seek to find

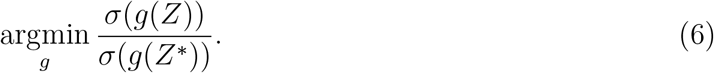

A prohibitive pitfall of this approach can come about by unconstrained shuffling of the data to produce *Z*^*^ in (6). In fact, the CR can be brought to its minimal value of zero, if there is a property that is always satisfied by the data but is violated by the shuffles (see the supplementary material Section 7.1, (25) for a proof). With such a property, one can construct a combination which attains a value of zero on the data and nonzero on the shuffled data, rendering the CR zero. This solution of the simple optimization problem holds information about the geometric distribution of data points but does not necessarily identify a regulated combination.

We illustrate this for the protein example presented above, where the ratio 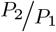 is maintained around a stable setpoint. The average CR of this combination over 100 realizations is not zero but rather 0.17 ± 0.09 because of random fluctuations. Conversely, a combination with a zero CR can be constructed based on the following general property of the data: in the observed time-series, 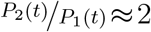 with small noise such that the data always satisfies *P*_2_ > *P*_1_. This constraint is not obeyed by the shuffled data: some values of *P*_2_ are smaller than *P*_1_ at other time-points, so that some shuffled traces will have 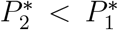. Thus, for example, the combination *g*(*P*_1_, *P*_2_, *P*_3_) =sgn(*P*_2_ − *P*_1_) is 1 at each time point yielding a standard deviation of zero for the real data, but different from zero in the shuffled data. Consequently, the CR is equal to zero. This artefact creates a potential pitfall to a simple optimization of (6).

Let us demonstrate this effect by a specific implementation, where the CR is optimized by an artificial neural network. Stretches of data similar to those presented in Figure 1A are fed into the network, with the target of minimizing the CR. Using a nonlinear network allows to search for combinations which are not necessarily linear, such as the desired ratio or the undesired sign function in this example. The network provided as output an optimal combination *g*(*P*_1_, *P*_2_, *P*_3_) that can be computed for any value of *P*_1_, *P*_2_, *P*_3_ but, due to the nonlinearity, cannot be easily expressed as a simple analytic formula.

To gain intuition into the combination found by the network, we compute *g*(*P*_1_, *P*_2_, *P*_3_) over the shuffled ensemble, and plot its value as a function of *P*_1_ and *P*_2_. Figure 2 depicts this value as a colormap in the (*P*_1_, *P*_2_) plane. Our prior knowledge of the true regulated combination in this example, allows us to mark its value (red line, depicting 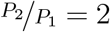), and moreover to delineate the region where real measurements occur (two white lines). Examination of this figure reveals that the optimal combination found by the neural network is practically constant on shuffles that remain within the limits of the data (flat light green area between the white lines). Outside these limits, it obtains varying values correlated with the distance of the shuffled point from the real data.

**Figure 2:**
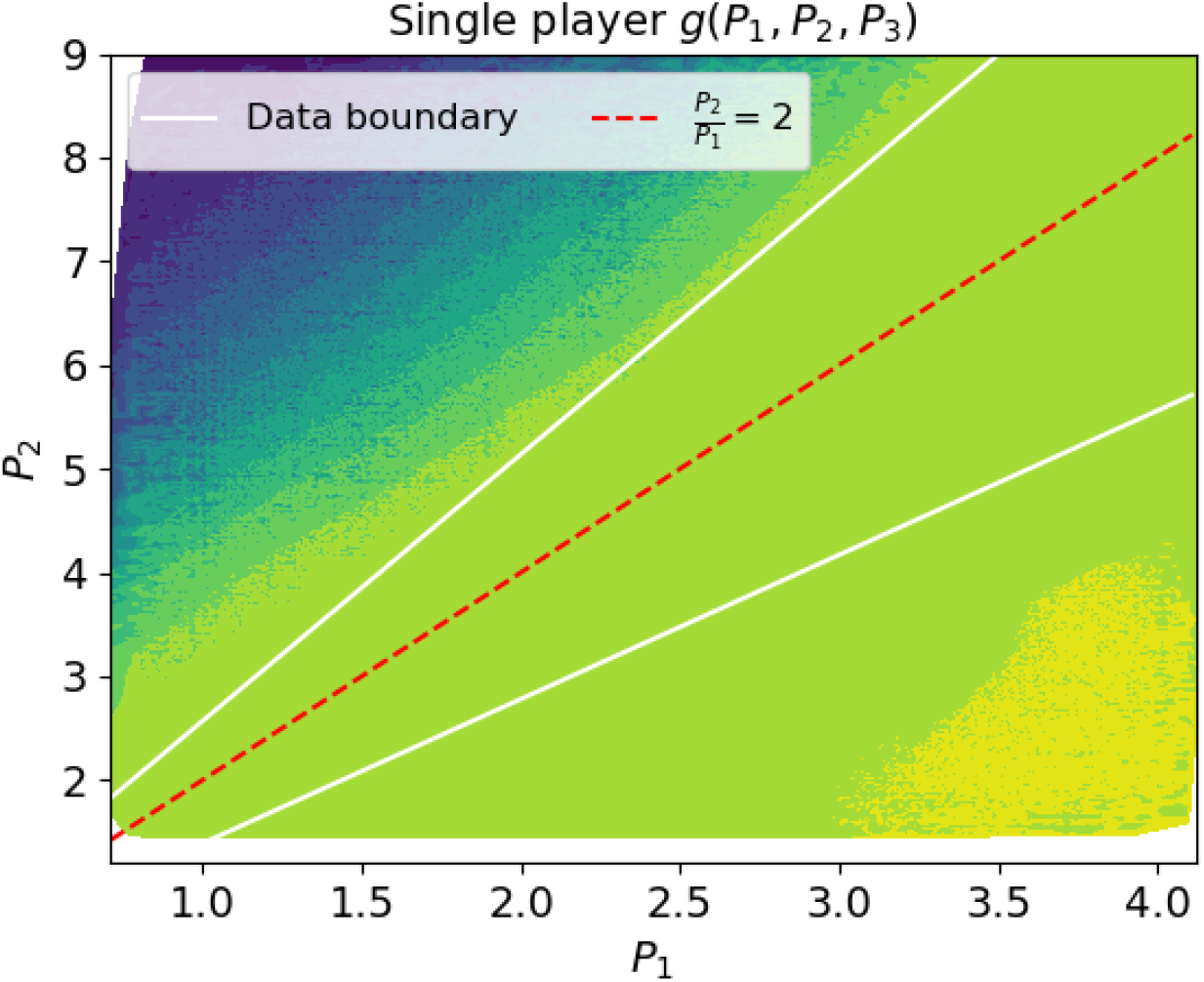
Failure of straightforward optimization. Optimal combination values found by the single-player algorithm, a neural network which minimizes the Coefficient of Regulation with unconstrained shuffling (CR, (5), *ζ*(·) ≡ 1). This algorithm was fed with time traces of the three proteins, with the ratio 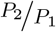 being the conserved combination. The network output is displayed as an arbitrary-value colormap in the (*P*_1_, *P*_2_) plane. Shuffles that fall within the boundaries of the data (the white lines) have a practically fixed value, while shuffles outside these boundaries attain values that are correlated with their distance from the boundaries.The found combination has a CRnof almost zero (0.004 ± 0.003). However, its Pearson correlation with the ground truth conserved combination is 0.11 ± 0.08.

This suggests that the optimal combination found by the neural network identifies the region occupied by the data, presumably by constructing an indicator function as described qualitatively above. Indeed, the output combination has a CR value of nearly zero, but a very low correlation with the true regulated quantity 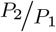. We refer to this algorithm as the “single player” since it involves only a single optimization goal: a “combination player” that aims to find a combination minimizing the CR. Analyzing its failure allows us to identify a way to correct it: if we constrain the shuffles to a set that is plausible in light of the data distribution, we may prevent the optimization algorithm from constructing artefact functions that reflect structural differences between the measured and shuffled ensembles. This will hopefully push the algorithm to discover the true regulated combinations, which reflect coupling between variables. We therefore introduce into the game a second player, whose goal is to constrain the shuffle ensemble such that it better resembles the data distribution, while still destroying temporal relations. This will be called IRAS (the “two player algorithm”) and will be described next.

### 2.3 IRAS captures the control objective

In the previous section, we reasoned that constraining the shuffled ensemble to be more similar to the real data may avoid artefacts and lead to meaningful combinations. Inspired by the concept of two competing players as implemented in the Generative Adversarial Nets algorithm [Goodfellow et al., 2014], we developed a scheme that alternates between optimizing the CR and constraining the shuffled ensemble.

The first player, termed the “combination player”, is a neural network that takes a step towards minimizing the CR. Following this step an adversarial player, denoted the “shuffle player”, resamples the shuffle ensemble to mimic the statistical structure of the data. This is done by resampling the shuffle ensemble such that the distribution of the current combination matches that of the original data. At the end of this step, the value of the CR equals 1, and the combination player starts another round of optimization, with the new resampled shuffled ensemble. We refer to this as IRAS, the two-player algorithm (see Figure 3). A detailed description of the resampling procedure is given in the supplementary material Section 7.2.1.

**Figure 3:**
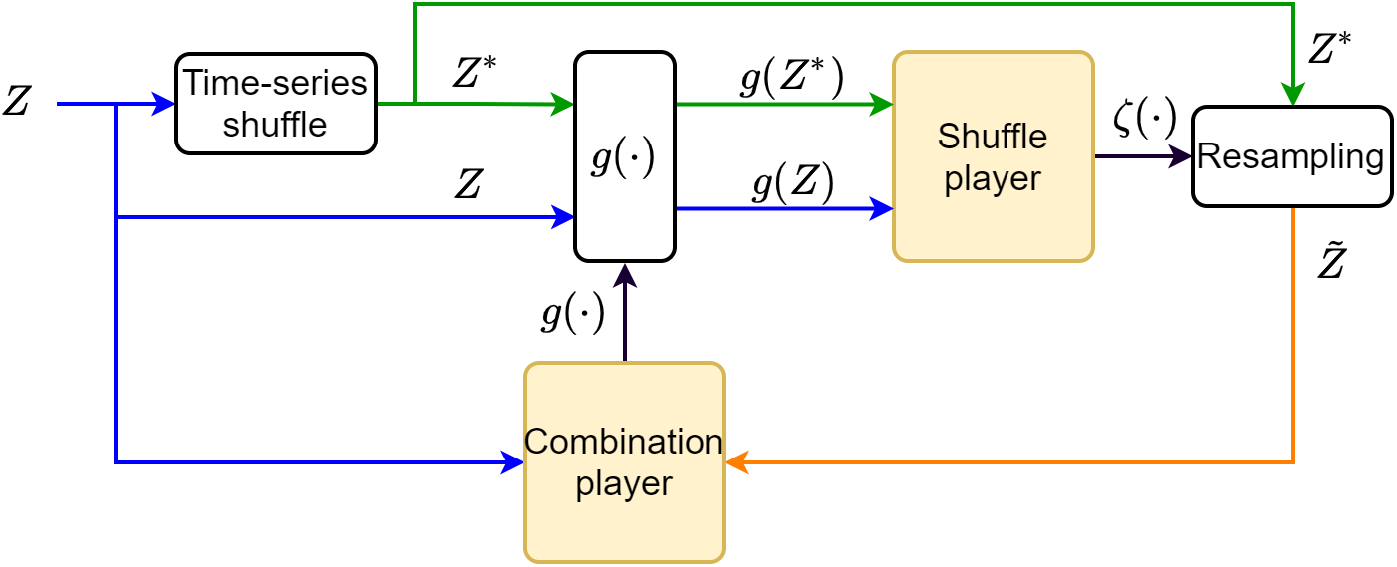
IRAS Algorithm outline. The time-series data *Z* is shuffled to create the unconstrained shuffled time-series *Z*^*^. The “shuffle player”, exposed only to the 1D combinations *g*(*Z*) and *g*(*Z*^*^), sets the weighting function *ζ*(·) used to resample 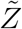 from *Z*^*^, such that the 1D distributions, *P*_*Z*_(*g*(*Z*)) and 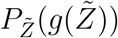, are identical. Then, given *Z* and 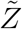, the “combination player” updates *g*(·) towards minimizing its CR. These steps continue to iterate until no further improvement is possible.

To demonstrate IRAS in action, we return to the protein example. Figure 4A depicts the progression of steps of the two players at a stage of the iterative optimization. A colormap of the combination in the (*P*_1_, *P*_2_) plane, found by the combination player, is shown on the top left. This combination defines a probability distribution on the real and shuffled data (top right). The shuffle player constructs a new shuffled ensemble by resampling (bottom right); by construction, in this new shuffle ensemble the distribution of *g* matches that of the data (bottom left). The combination player receives this updated shuffled ensemble and the next optimization step begins.

**Figure 4:**
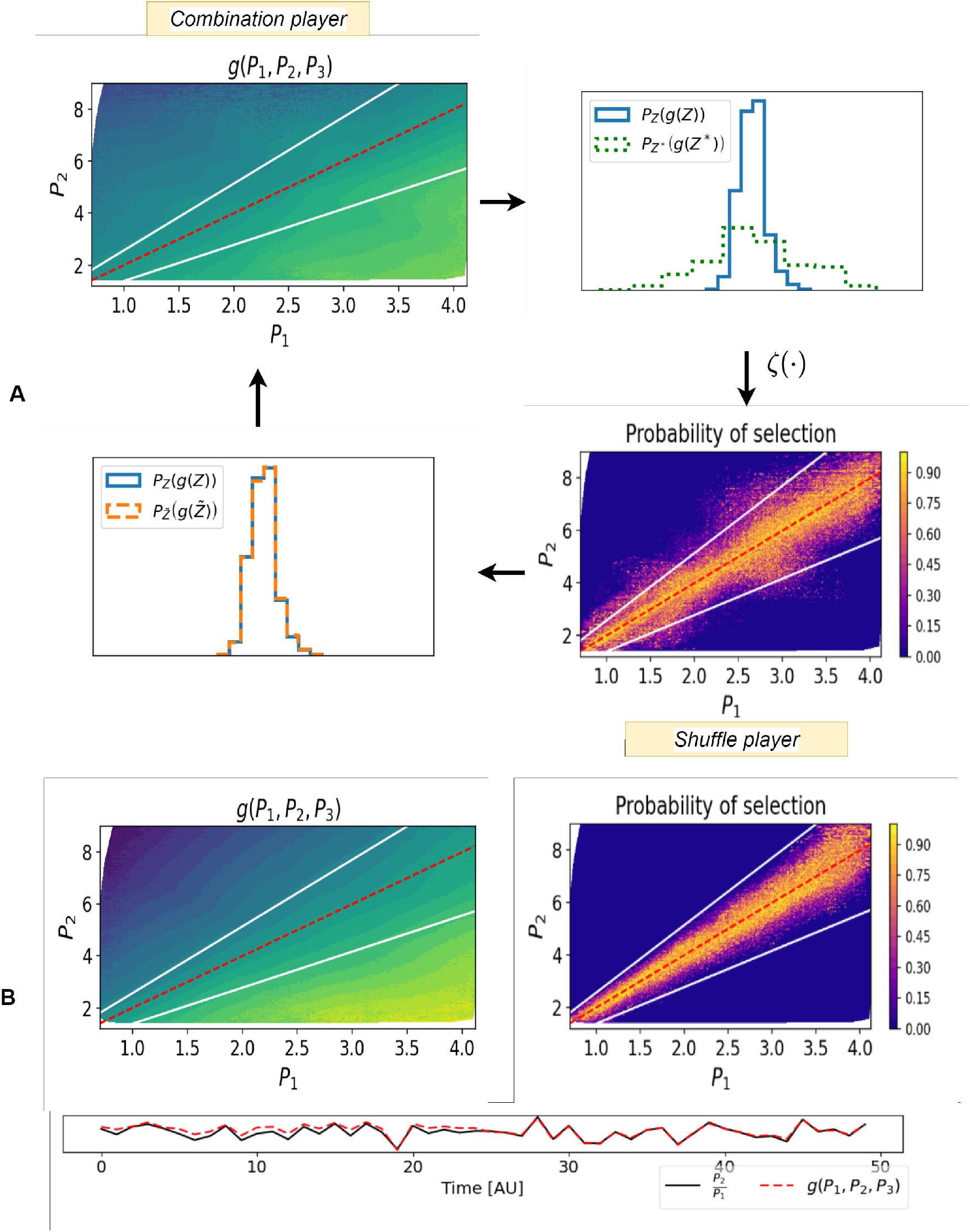
IRAS demonstration. (A) A step of the iterative algorithm (see Figure 3) is displayed. Top left: the value of an intermediate combination *g*(·) given by the combination player, is displayed in the (*P*_1_, *P*_2_) plane together with the true combination (red line) and the data limits (white lines). Top right: distriubtions of this combination over the data (*P*_*Z*_ (*g*(*Z*)), blue) and unconstrained shuffles 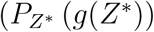, dashed green). Bottom right: The shuffle player examines these 1D distributions, and resamples *Z*^*^ via the weighting function *ζ*(·) to construct the constrained shuffles 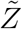, over which the 1D distribution of *g* matches the data. The resampling probability is displayed in the (*P*_1_, *P*_2_) plane. Bottom left: Combination player receives this resampled shuffle ensemble, another optimization step begins and the combination player updated *g*(·). (B) Combination values (top left) and resample probability (top right) at the final iteration. The combination player has captured the control objective (the map approximates 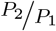), and the shuffle player has captured the data distribution (delineated by white lines). Bottom: values of the combination along a stretch of time together with the ground-truth combination. 11

Gradually, the resampled shuffle ensemble approximates the distribution of the real data and a map which approximates 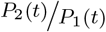 emerges as the output combination. At the final step, Figure 4B, the combination player cannot further minimize the CR. We find that indeed IRAS converges and outputs the true conserved quantity, 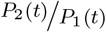, as observed in the bottom row of Figure 4B^*^.

## 3 Validation

After presenting the construction of IRAS, we seek to validate it on datasets with a known control objective or other conserved quantity. We chose two validation examples from different biological scales, a kinetic model of protein interactions and a measured dataset from a psychophysical experiment. Additionally, to demonstrate the efficiency of our algorithm in studying mechanical physical systems we model a simple physical spring system where energy is conserved. Throughout the validation examples we use a single neural network architecture whose details are listed in the Supplementary material Section 8.

### 3.1 A kinetic model of regulatory interactions

We first validate IRAS on simulated data generated from a kinetic model that describes regulatory interactions between three proteins incorporating a feedback loop. In the considered model (inspired by [El-Samad, 2021]), the total amount of two proteins *P* and *S*, namely *P* +*S*, is controlled by another protein *M* under perturbations in protein expression parameters. The model (see illustration in Figure 5A) is described by the differential equations

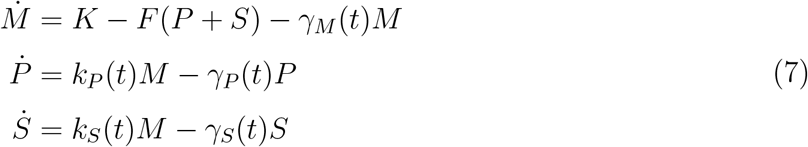

where the three proteins *M, P, S* are linked in a feedback loop. Both *P* and *S* are positively affected by *M*, with their steady-state values proportional to it. The concentration *M* in turn, is negatively affected by the sum *P* + *S*, with the strength of this negative feedback given by the rate constant *F*. The degradation rates *γ*_*M*_, *γ*_*P*_, *γ*_*S*_ and the production rates *k*_*P*_, *k*_*S*_ are perturbed over time as shown in Figure 5B (top panel). Figure 5B (bottom right) shows the trajectories of the three proteins across time. Small changes in *S* or *P* induce swift and sharp changes in the production rate of *M* and maintain *P* +*S* around a stable level. This is reflected in a high negative correlation between *S* and *P*.

**Figure 5:**
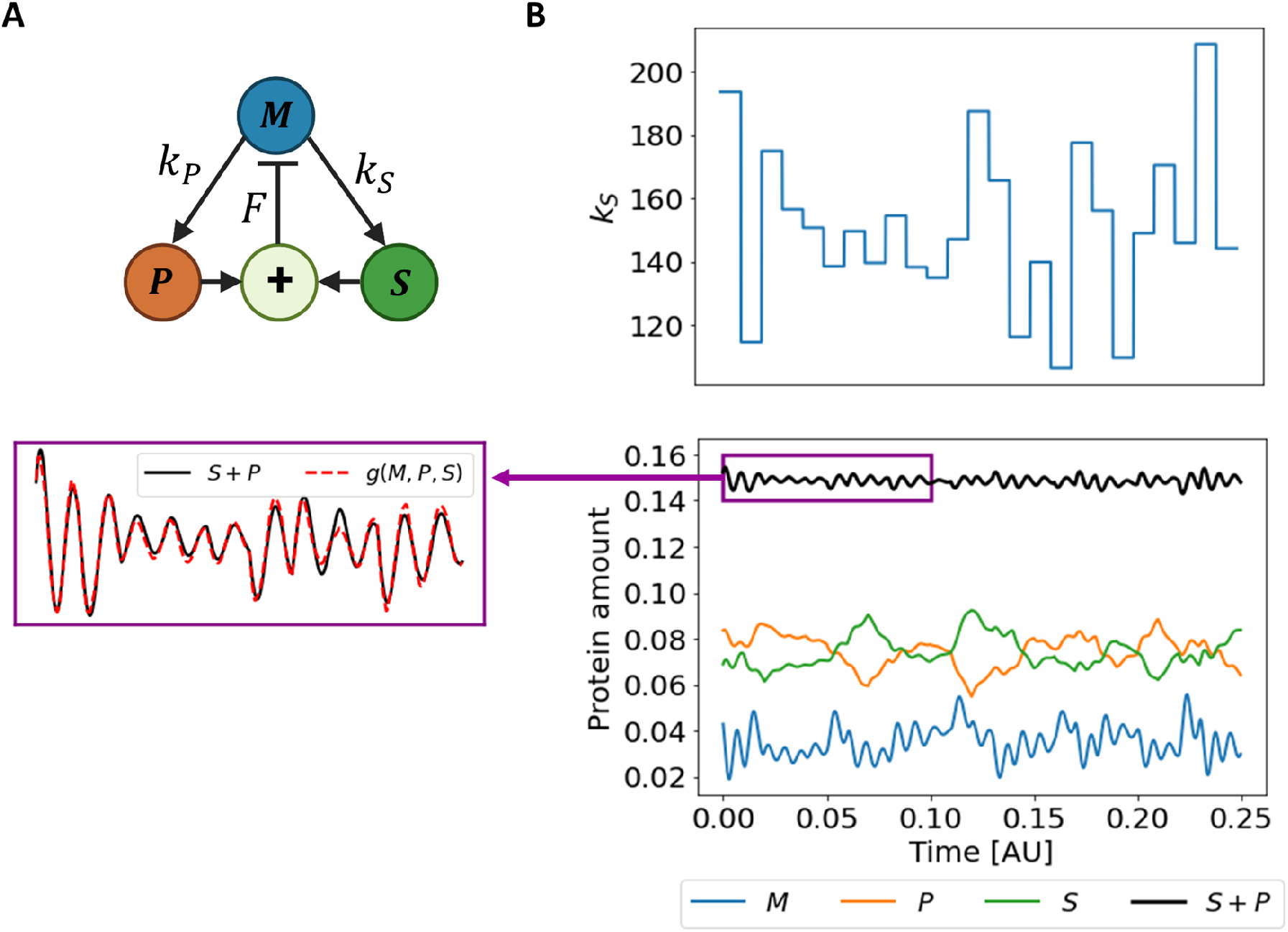
IRAS captures the control objective in a kinetic model. **(A)** An illustration of the closed loop system. Protein *M* induces the production of both *S* and *P* and receives a negative feedback of their sum. **(B)** Top panel: perturbations cause step-like variation in *k*_*S*_ over time. The duration of each step is 0.01 which is much longer than the time scale of the feedback loop *τ* = ^1^*/*_*F*_ = 0.0005 (*F* = 2000). This enables the controller to track the changes in *S* + *P*. Each step was sampled from a normal distribution with a mean 150 and a standard deviation 30. The rest of the parameters were sampled similarly: *γ*_*P*_, *γ*_*S*_ = 70±15, *γ*_*M*_ = 80±15, *k*_*P*_ = 150±30. Bottom right: The trajectories of the three proteins and the combination *S* + *P* over time. The Bottom left: A zoom-in of the combination *P* + *S* (black) within the purple box in the right panel along with the output of the algorithm (dashed red).

To gain a better insight into the stability of the combination *P* +*S*, we consider the steady state of the system for a single constant parameter set. At steady state, the rate of change of all three proteins is zero, and *P* +*S* is given by

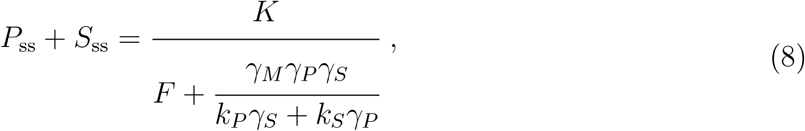

where *P*_ss_ and *S*_ss_ are the steady states of *P* and *S* respectively (see supplementary section 10.1 for the steady states of the three proteins). If the strength of the negative feedback is large and satisfies 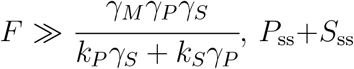 will approximately remain around the same setpoint despite the perturbations. This indicates that *g*(*M, P, S*) = *S*+*P* is a possible control objective of the system under these conditions. Indeed, applying IRAS on 30 realization of this model, we find that it outputs the control objective *S* + *P* with high accuracy; Figure 5B (bottom left) shows their overlap. The mean Pearson correlation between them is 0.97 ± 0.005 over the 30 realizations. We present in Supplementary Information section 10.2 additional examples, including cases where different parameters lead to different conserved combinations, that are still captured by the algorithm.

In summary, IRAS identifies correctly the control objective in a dataset generated from a kinetic model of regulatory interactions.

### 3.2 Relational dynamics in perception

Human perception is inherently noisy and the source of this noise is an important issue in Psychophysics research [Faisal et al., 2008, Monto et al., 2008, Marom, 2010]. An experimental design was introduced to address this question, which involves a closed-loop controller that modulates the input stimulus according to the human responses, with the goal of decreasing variability and maintaining the response probability at a pre-determined setpoint [Marom and Wallach, 2011], see Figure 6A. It was shown that this feedback loop indeed quenches the response variability.

**Figure 6:**
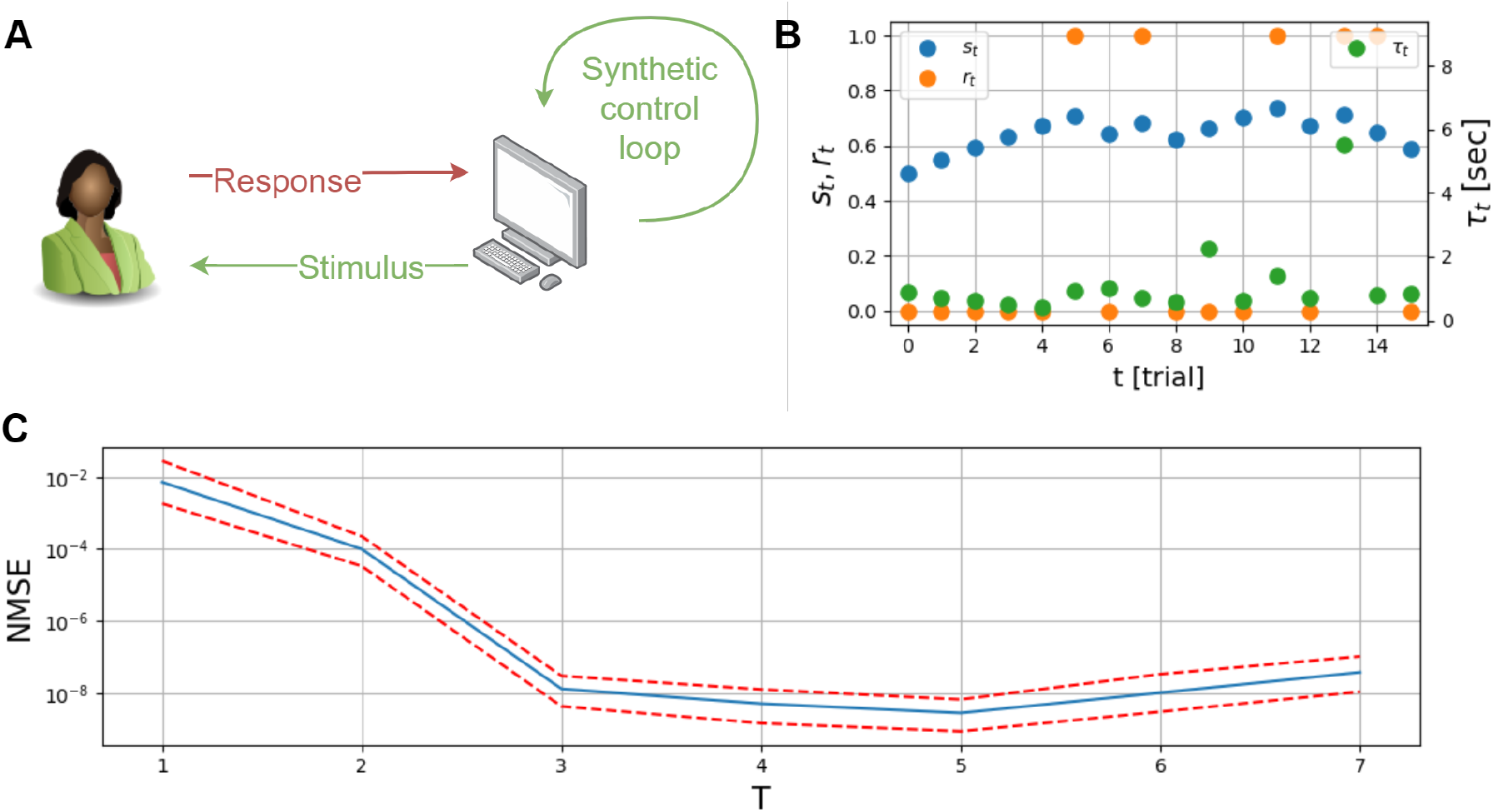
Relational dynamics in perception. (A) Trial-trial variability in human sensory detection is tested. A synthetic feedback controller sets the stimulus, which is the contrast of a foreground raster displayed on the screen. Then, the user responds positively when detecting the raster or negatively when not. (B) Raw data from psychophysics experiment. A portion of the measured time series *z*_*t*_ = [*s*_*t*_ *r*_*t*_ *τ*_*t*_] in a human sensory detection experiment. The stimulus *s*_*t*_ (blue dots) is a time-series of image-contrast values, the responses *r*_*t*_ (orange dots) are a Boolean time-series of detection, and *τ*_*t*_ (green dots) is the reaction time from stimulus to response. (C) Normalized mean-square-errors of stimuli estimation values as defined in (14). The stimuli estimation is obtained from the analytical expression of the feedback loop detected. The estimation errors decrease monotonously up to *T* = 5 implying an effective time-scale of 5 trials. Dashed red lines are the MSE of errors higher and lower than the MSE which lies on the blue line.

The data from these experiments provide a unique opportunity to apply and validate IRAS, as we have a ground truth component in the system - the engineered controller that records the human responses and determines the next stimulus. This controller is coupled to a noisy biological system, the human observer. For validating our algorithm, we feed it with the complete raw data, including both input stimuli and responses. If working correctly, IRAS should identify the synthetic controller as the most regulated combination and allow us to derive a mathematical description for the way it sets the stimulus. We emphasize that the algorithm does not have access to any internal variables of the synthetic controller.

The task in the experiment of [Marom and Wallach, 2011] consisted of sensory detection of a weak visual stimulus. In sequential trials, users were presented with a random raster of black and white pixels. A smaller foreground raster drawn from a different distribution, was embedded in the background raster area. A single session was composed of multiple trials, where the foreground raster was displayed at a random location on the screen; in some of the trials only the background was displayed. In each trial, users had to respond if they detected the foreground raster and withold response if not. The synthetic feedback controller set the contrast of the foreground raster as a function of the previously received responses, increasing it when response probability was low and vice versa, with the objective of maintaining a fixed probability of response. The response time was also recorded in each trial, but the controller did not make any use of this information.

The experiment was performed on eight human subjects yielding a dataset that consists of three-dimensional, discrete time-series, including the stimuli, responses and reaction times, over 450 trials for each subject. Figure 6B depicts a portion of the three components of raw data as a function of trial number *t*. The three observables are the raster contrast levels (*s*_*t*_ ∈ ℝ, blue); corresponding binary responses (*r*_*t*_ ∈ {0, 1}, orange); and reaction times (*τ*_*t*_ ∈ ℝ, green). Our validation here will consist of feeding this data to IRAS to find the most regulated combination.

Recall that in Section 2, IRAS was presented for the case where the most regulated combination of measurements is sought among instantaneous functions *c*(*t*) = *g*(*z*(*t*)). The shuffle player created an ensemble where all correlations among observables measured at the same time point were eliminated. Here we would like to derive the most regulated combination between measurements at consecutive time points. We expect that this will allow for the identification of the synthetic controller that sets the stimulus *s*_*t*_ as a function of past values. To this end, the shuffled data provides a random *present* for a given *past*, while preserving correlations within the same time point - and thus preventing the algorithm from detecting a combination that does not relate past and present observations. Over *T* + 1 consecutive observations (*T* > 0), we now seek the most regulated combination constructed as

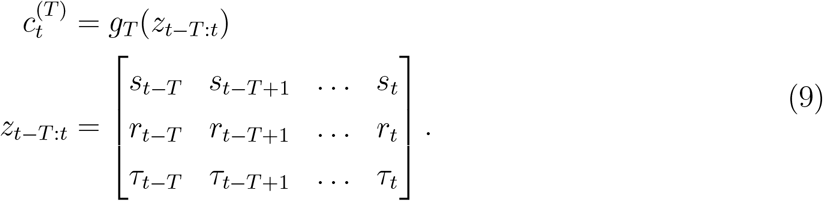

While the shuffle player creates surrogate data of the form

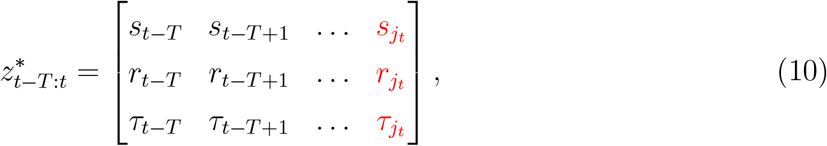

where 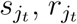 and 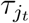 are the observations obtained at some random time *j*_*t*_. We refer the reader to the supplementary material Section 7 for the complete technical details of the implementing IRAS over varying time-windows of size *T*.

We ran the algorithm on the experimental dataset with this definition of shuffles, for different values of *T*. In each evaluation, the yielded combination *g*_*T*_ (·) is the output of an artificial neural network, therefore it is effectively a black-box. In this case, we could approximate the network by a multivariate polynomial and obtain an interpretable expression while remaining close to the actual network output (Pearson correlation of 0.88 ± 3*e*^−5^ over the 8 human subjects). For *T* = 3, the resulting approximation is,

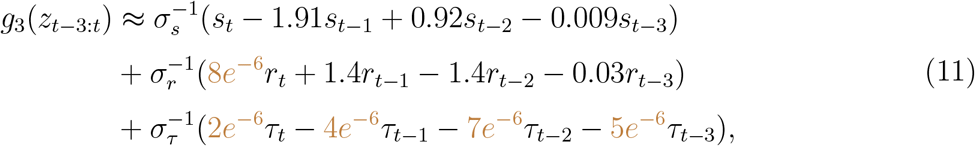

where

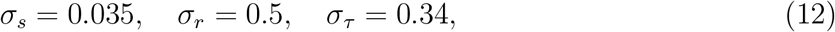

are the standard deviations of the stimulus, the response and the reaction time respectively.

Comparing these results to our prior knowledge of the experimental system provides strong support to the validation of IRAS. We expect the synthetic controller to be identified by the algorithm, and therefore that the current stimulus will be a function of previous responses. This should translate to small coefficients for *r*_*t*_ and *τ*_*t*_, compared to that of *s*_*t*_. Furthermore, all reaction time coefficients should be negligible, because they were not used by the controller. Examining the coefficients in (11) shows that these are indeed properties of the discovered combination. To examine whether this combination captures the synthetic feedback control loop, we test whether the obtained combination can predict the stimulus values correctly. Removing the negligible terms in (11) and recalling that *g*_*T*_ (·) ≈ *c*_set_ we predict (up to a constant term),

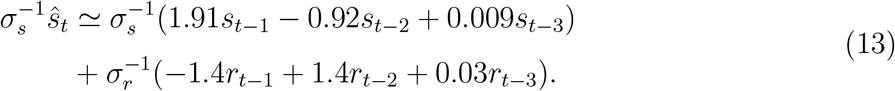

Here we denoted the stimulus obtained from the learned combination by *ŝ*_*t*_ so that it can be compared to the true stimulus value *s*_*t*_, set by the controller in the experiment. The normalized mean-square prediction error,

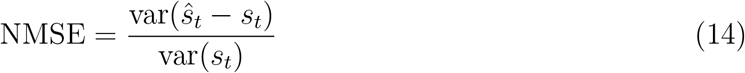

is extremely low, approximately 10^−8^, testifying to a high degree of functionality of the internal synthetic feedback loop detected by IRAS.

Another important question that the our methodology allows to address, is the effective time-scale of the feedback loop. Running the algorithm on various values of *T*, we estimated the mean-square prediction error (14). Figures 6C and 15 show the result as a function of *T*, testifying to an effective time-scale of 5 trials. Because the system works in closed-loop, this timescale cannot be directly compared to the controller timescale.

In summary, IRAS identifies correctly the most regulated combination, corresponding to a synthetic control feedback loop, in data obtained from a real-world experiment with a human in the loop.

### 3.3 Identifying conservation laws

IRAS identifies quantities that are maintained at approximately constant values throughout the dynamics, based on the empirical criterion of CR. There can be different underlying reasons why a quantity remains fixed. For biological systems, such behavior may indicate regulation - i.e. compensation that protects some variables from perturbations. In physical systems, constant values often represent exact conservation laws.

In the previous sections we validated IRAS on datasets of simulated and experimental biological systems. The assumption was that different realizations of the simulation, or different experiments, were statistically similar. Specifically, we assumed that all systems share the exact same *g* function. More generally, different systems might share the same functional form for *g*, but with different system-specific parameters. In this section, we demonstrate how IRAS deals with a family of datasets that stem from systems with different parameters. We do this in the context of a physical system, illustrating the wide applicability of IRAS. As an example for this challenge, we focus on a Hamiltonian mechanical system [Reichl, 1999, Sakurai and Commins, 1995]. Hamiltonian mechanics is a branch of physics that describes dynamical systems through conservation laws and invariances. The Hamilton-Jacobi equations relate the state of a system to some conserved quantity, e.g. energy. In physics, specific a-priori knowledge of the system is required to identify its invariants; finding invariants in a general dynamical system, or even knowing whether or not they exist, is a difficult problem. Developing automated computational methods to find invariants from data is a challenge of much recent interest [Watters et al., 2017, Santoro et al., 2017, Hamrick et al., 2018, de Avila Belbute-Peres et al., 2018, Chang et al., 2016, Tenenbaum et al., 2000]. A conservation law is a function that satisfies (1), therefore IRAS is suitable for its identification. We note that optimizing for conservation alone can lead to trivial quantities, such as predicting a constant *g*(*z*) = *c* independent of *z*. In a recent paper, [Alet et al., 2021] refer to a non-trivial *g*(·) by the term *useful conservation law*.

We consider the ideal frictionless mass-spring system shown in the upper right corner of Figure 7A. This system is commonly used to test algorithms designed for the identification of conservation laws [Greydanus et al., 2019, Alet et al., 2021]. The system’s Hamiltonian and dynamic equations are,

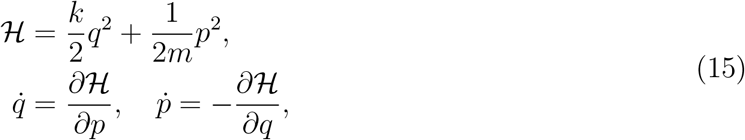

with two parameters: the spring constant *k* and the mass *m*. The dynamic variables are *q*, the coordinate denoting deviation from equilibrium, and the momentum *p*. The most conserved instantaneous combination is the Hamiltonian, reflecting conservation of energy. We examine several systems with different parameters, such that the value of the conserved quantity differs between them but the functional form is the same. The simulated dataset contains 100 different mass-spring systems. In each system, *k* and *m* were sampled uniformly between [0.5, 1.5] and the initial conditions, *p*_0_ and *q*_0_, were sampled uniformly between [0.15, 0.25] and [0.1, 0.2] respectively. The raw observations consists of times-series of length 1000 of *p* and *q* corrupted by a zero-mean additive Gaussian white noise with standard deviation 0.01. Figure 7A shows the observed traces as a function of time (left) and in the (*p, q*) phase plane (bottom right) for three systems. Figure 7B, left, shows the corresponding time-series of the Hamiltonian, namely the energy as a function of time.

**Figure 7:**
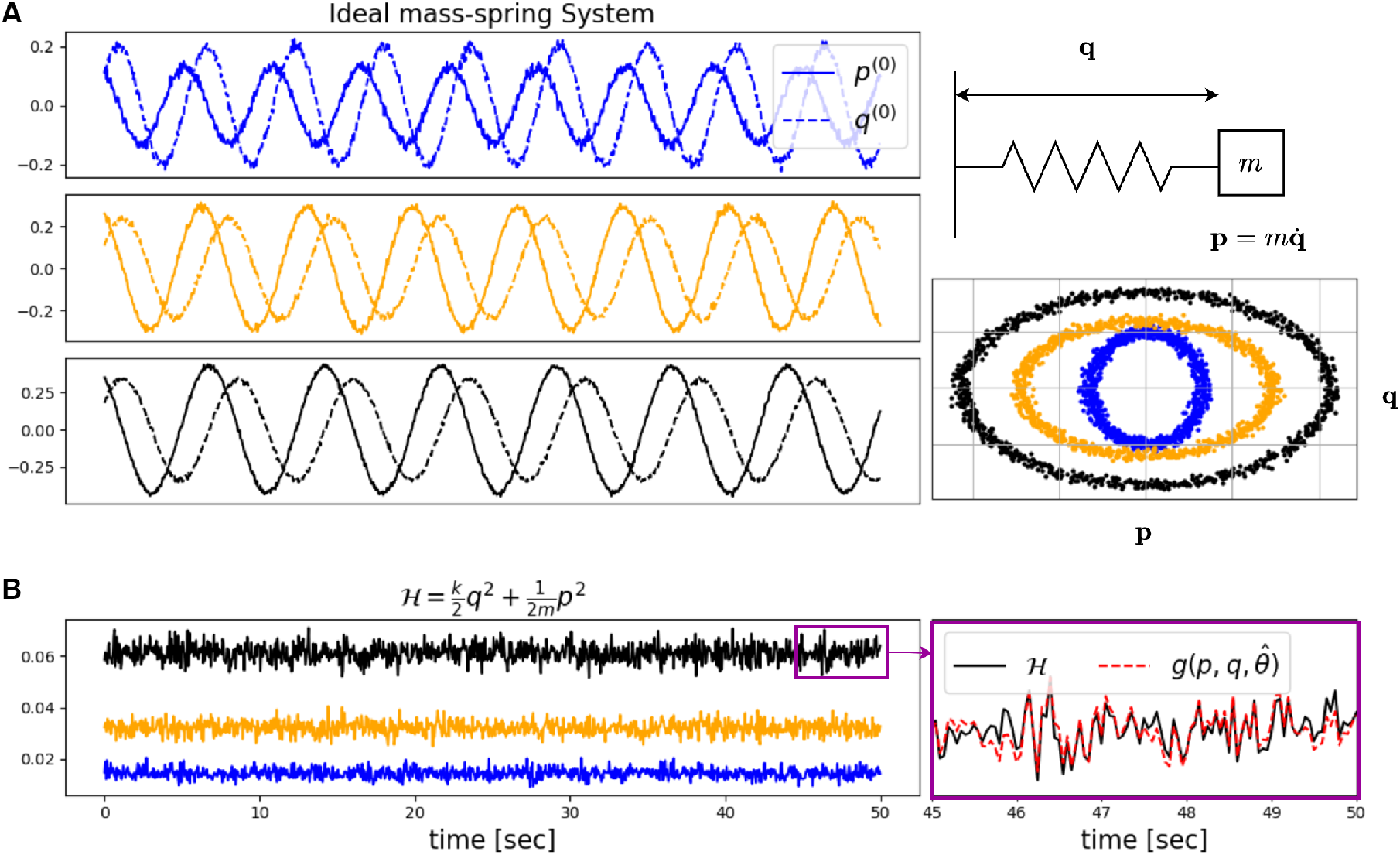
IRAS captures the conservation law in Hamiltonian mechanics. (A) Top right: ideal particle on spring with mass *m* and spring constant *k*. Left: Time traces of momentum, (*p*, solid lines) and coordinate, (*q*, dashed lines) for ideal mass-spring systems with spring and mass constants *k* = (0.72, 1.13, 1.05) and *m* = (0.59, 1.32, 1.48) in the top, middle and bottom panels respectively. Bottom right: same trajectories plotted in the phase plane (*p, q*). (B) Left: The energy as a function of time for the three systems in (A) with corresponding colors. Right: zoom of the energy and output of the combination learned by IRAS in a short stretch of time.

The different physical systems share the same conservation law with the Hamiltonian as the invariant combination. However, the value of this combination is different in each system and depends on the parameters. Running IRAS over the pooled data from all systems, in the same setting used in the previous sections, that is, optimizing a combination *c*(*t*) = *g* (*p*(*t*), *q*(*t*)), leads to a low mean Pearson correlation of 0.65 ± 0.15. This low value occurs because the learned combination *g*() did not incorporate system-specific parameters. To address this problem, we now present an extension to IRAS that allows for identifying a regulated combination that is a function of both the measurements and of parameters that are estimated simultaneously for each system (detailed in the supplementary Section 7). In the extended version, the learned instantaneous regulated combination is,

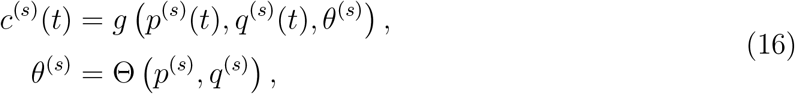

where super-script *s* is for system *s* ∈ {1, 2, …, 100}, and where *p*^(*s*)^ and *q*^(*s*)^ are the timeseries observed from system *s*. The set of parameters estimated for system *s* is *θ*^(*s*)^ ∈ ℝ^*l*^ with *l* a user-defined hyper-parameter, and where Θ(·) is a second artificial neural-network (see Figure 12 in Supplementary Section 7). Here we set *l* = 2. Indeed, the extended IRAS captures the conservation law yielding a mean Pearson correlation of 0.95 ± 0.012 between *c*^(*s*)^ and ℋ^(*s*)^ (averaged over all systems). Figure 7B, right, shows the values of ground-truth ℋ together with the learned combination *c* along a stretch of time for a single system. The parameters estimated by Θ(·) match the physical quantities of spring and mass constants, exhibiting Pearson correlation of 0.88 and 0.82 with 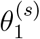 and 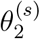respectively (averaged over all systems).

As a limiting case we tested the extended algorithm, (16), in a scenario where all mass-spring systems are identical with *k* = 1 and *m* = 1. The resulting Pearson correlation between ℋ and the identified combination is 0.96 ± 0.01. This testifies that the extended algorithm does not negatively affect the performance when estimating system-specific parameters is unnecessary.

In summary, IRAS correctly identifies a conservation law which is the most “regulated” (conserved) combination in a Hamiltonian mechanical system. With the extension of estimating a user-specified number *l* of parameters for different systems, the algorithm can identify the relevant physical parameters and construct the conserved quantity as a combination of the instantaneous observations and the estimated physical parameters.

## 4 Discussion

Detecting invariants in dynamic data is a technically challenging problem with many potential applications. We presented IRAS, an algorithm that receives as input raw dynamic measurements and provides combinations, or functions, of these variables that are maximally conserved across time. Such conservation can be the result of feedback control, where some variables compensate others to protect a control objective; or it can be the result of symmetry and exact conservation laws. Taking a phenomenological approach to the problem, we introduced a quantitative measure – the Coefficient of Regulation (CR) – that characterizes the sensitivity of a combination to destroying temporal order among its constituents. This measure, regardless of the mechanism underlying invariance, serves as the basis of an optimization algorithm that outputs the combination most severely affected by temporal shuffling.

While the CR is an intuitive measure, and can be shown to be very small for regulated or conserved combinations, its straightforward optimization is insufficient to escape “trivial” combinations that do not provide meaningful relations between the variables (see Section 2.2). Related problems were highlighted in previous work, where trivial constants resulted when attempting to identify conserved quantities in physical systems [Alet et al., 2021]. To identify meaningful combinations with small CR, some constraints on the shuffle ensemble need to be implemented, so that time-ordering is destroyed while respecting the boundaries of the data. We proposed an iterative process between two players, one minimizing the CR and the other creating successively more constrained shuffle ensembles. IRAS converges when the two players cannot further improve. The neural network then outputs a combination of variables as a proposed conserved quantity.

We provide validation in three distinct examples taken from very different realms, and which reveal three versions of IRAS. First, we validated the simplest version, which optimizes an instantaneous combination – namely, a function of the dynamic measurements at the same time point. Using a kinetic model of interactions between proteins, with feedback regulating a sum of two of them, we simulated traces over time in which system parameters were randomly perturbed. The feedback in the system induced compensations that maintained the control objective at a setpoint. This objective was correctly identified by the algorithm.

Second, we analyzed data from a human-computer closed-loop visual detection experiment [Marom and Wallach, 2011], where the computer implemented a feedback loop that clamps the human response. Here, the CR was minimized among combinations that include consecutive time points in the data, aiming to recover the target of the engineered control system. The qualitative dependence between variables was identified correctly.

Finally, we investigated a physical system with a symmetry-related conservation law. Here we presented a generalized version of IRAS, suitable for cases where data from many similar systems are available, each with a different parameter set. Another neural network was added which identifies the parameters simultaneously with the two-player CR optimization (see Figure 12). We emphasize that also here, no prior knowledge regarding the physical parameters was used. We demonstrated the success of the algorithm in identifying correctly the energy constant in a collection of ideal spring systems with different masses and spring constants, in the presence of noise. This result is significant in light of the difficulty to identify conserved physical quantities, even in a single system [Greydanus et al., 2019].

The conserved quantity is an implicit relation between measurements, defining a constraint and thus effectively reducing the dimensionality of the data. This is somewhat reminiscent of dimensionality reduction problems. The goal, however, is quite distinct in the two scenarios. Dimensionality reduction aims to describe the maximal amount of variability in the data using as few descriptors as possible. In contrast, we aim to find a **meaningful** combination of the data with a minimal amount of variability. The restriction to meaningful combinations, achieved through temporal shuffling, renders the two approaches qualitatively different and not easily comparable.

Our main motivation in this work was to understand regulation in complex biological systems. Often, such systems are “reverse engineered” – for example, using System Identification or other methods, to build mathematical models based on observed data [Ljung, 1998, Schmidt and Lipson, 2009, Chen et al., 2022, Daniels and Nemenman, 2015, Shen et al., 2021, Haber and Schneidman, 2022]. A model can then be investigated to shed light on the functionality of the system, its robustness and other properties. However, multi-variable, multi-parameter models are generally hard to understand even with explicit equations; parameter space can be vast, different parameters can lead to markedly different behaviors, and it is not always clear what the system’s functionality is.

Instead, we propose to analyze the properties of complex biological systems *bypassing* the modeling stage, to provide insight directly from dynamic data. We ask the general question: “what does the system care about”? in the sense of control theory. Namely, we seek to identify, directly from data, conserved or regulated quantities that the system protects from fluctuations and perturbations. Such homeostasis is a phenomenon of central importance in many biological contexts.

Being a purely data driven analysis method, we tell the story of the system in the ‘language’ of the observables. Thus, we are limited by them. Spurious correlations between measurements may manifest as artefactual regulated combinations. Likewise, if by some fortunate coincidence, the controlled objective of the system is one of the individual raw measurements – IRAS will discard it, because it is not a combination. Both of these caveats highlight the importance of biological context in data analysis.

The presented algorithm detects the *most* regulated combination within the observables. Commonly, a biological system will regulate multiple different objectives via different feedback loops. Once the most regulated combination was identified we would like to continue the analysis and discover the next regulated combinations and be able to describe a hierarchy of control objectives [Stawsky et al., 2021]. This aim is left for future work.

## Acknowledgements

This work was partially supported by the Israel Science Foundation (grant numbers 451/17 (RM) and 155/18 (NB)), and by the Skillman chair in biomedical sciences (RM). We acknowledge the Adams Fellowship Program of the Israel Academy of Science and Humanities (AS). RM and RT are partially supported by the Ollendor Center of the Viterbi Faculty of Electrical and Computer Engineering at the Technion.

## Supplementary material

### 5 Feedback control in biological system models

Let the observed system be a non-linear dynamical system formulated by the stochastic differential state space model,

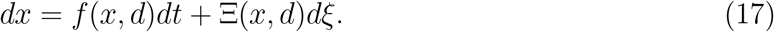

where both the deterministic term *f*: ℝ^*n*×*b*^ → ℝ^*n*^ and the stochastic term Ξ : ℝ^*n*×*b*^ → ℝ^*n*^ are a function of the state and an external input *d* ∈ ℝ^*b*^. The term *dξ* is an increment of a Wiener process. For details on the formalization of continuous-time stochastic differential equations we refer the reader to Chapter 3 in [Åström, 2012].

Biological systems internally regulate some of the state variables, or, more generally, some combinations of the state variables. We denote such an internally regulated combination by *c*(*t*) ∈ ℝ and denote by *c*_set_ the set-point value of the regulation such that the biological system maintains a small deviation from the set-point value, namely 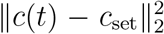 is small. Human-engineered systems are commonly represented by two separate entities – a plant and a controller. Biological systems, in contrast, are commonly modeled by a single dynamical equation. Therefore, the controlled objective and the control signal in a biological system model are implicit. Although control theory typically assumes the existence of a separate plant and controller (see Chapter 1.2 in [Åström and Murray, 2021]), most theoretical results and analysis tools, among them the analysis algorithm represented herein, do not require such a separation [Cosentino and Bates, 2011].

We do not observe the complete state of the system, but some function of the internal state that might be partial. Let *y* ∈ ℝ^*m*^ be a vector of observables which is some instantaneous function of the internal state, the input and a white measurement noise *v* ∈ ℝ^*m*^

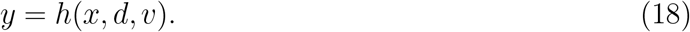

Since we only have access to the observations *y*, we seek some regulated combinations of observations, which, assuming observability^*^, should hold for the internal variables. In this study we present Identifying Regulation with Adverserial Surrogates (IRAS), a novel data analysis algorithm that receives a dataset of observed systems and yields regulated combinations that are jointly held by the observed system. These combinations are of two types: instantaneous combinations and combinations of measurements that are observed along a fixed-length time window.

### 6 Notations

#### Dynamical systems

- *x*^(*s*)^ ∈ ℝ^*n*^ - state of system *s*
- *θ*^(*s*)^ - parameter vector of system *s*
- *d*^(*s*)^ ∈ ℝ^*b*^ - external input to system *s*
- *f*_*s*_ : ℝ^*n*×*b*^ → ℝ^*n*^, Ξ_*s*_ ∈ ℝ^*n*×*n*^ → ℝ^*n*^ - functions describing the deterministic and stochastic components of system *s* such that *dx*^(*s*)^ = *f*_*s*_ (*x*^(*s*)^, *d*^(*s*)^) *dt* + Ξ_*s*_(*x, d*)*dξ*^(*s*)^ where *dt* is a time increment and *dξ*^(*s*)^ is the increment of a Weiner process.
- *y* ∈ ℝ^*m*^ - observation obtained from system *s*
- *v* ∈ ℝ^*m*^ - measurement noise in system *s*
- *h* : ℝ^*n*×*b*×*m*^ → ℝ^*m*^ - measurement equation of system *s* such that *y*^(*s*)^ = *h*(*x*^(*s*)^, *d*^(*s*)^, *v*^(*s*)^; *θ*_*s*_)

#### Dataset

- 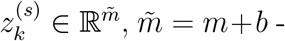 concatenation of the external known input *d*^(*s*)^ and the observation *y*^(*s*)^ sampled at time 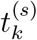
- 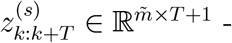 concatenation of *T* samples, 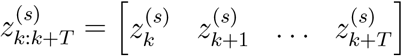
- 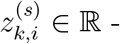 the i^th^ entry of 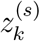
- *N*_*s*_ - number of samples 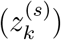 obtained from system *s*
- *M* - number of observed systems

#### Regulated signal and combination

- 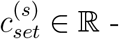 set-point value of system *s*
- 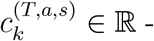 the value of the most *regulated signal* of system *s* at time *k* over a time-scale of *T* consecutive samples
- 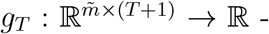 the most *regulated combination* over a time-scale of *T* consecutive samples for all *M* observed systems such that 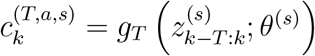

#### Algorithm variables

- Ω^(*T,α,s*)^ - a set that consists of *N* ^(*T,α,s*)^ members, *α* ∈ {*a, n, p*}
- 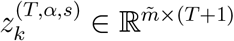 - a member of the set Ω ^(*T,α,s*)^
- 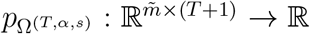 - the distribution from which members were sampled to the set Ω ^(*T,α,s*)^
- 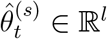 - estimated parameters of system *s* in iteration *t* of IRAS
- 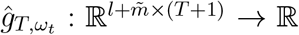 - the estimated *regulated combination*, a function parameterized by the parameters vector *ω*_*t*_
- 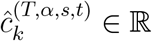 - estimated *regulated signal* of system *s* at time *k* in iteration *t* of Algorithm 1 such that 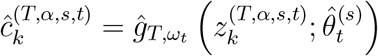
- 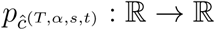 the distribution of *ĉ*^(*T,α,s,t*)^ such that 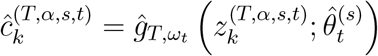.
- 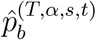 value of bin *b* of the histogram of 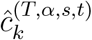 over all 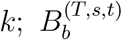set of values included in bin *b*

### 7 Problem formulation

As described in section 5, the systems we observe are non-linear stochastic dynamical systems. We analyze a dataset of observations of multiple similar systems, where the dynamic and observation equations of system *s* are given by

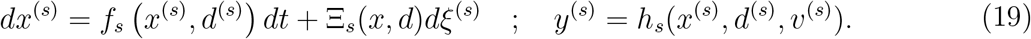

#### Discrete time formulation

Although we handle real-world systems that are by nature continuous time systems, we formulate the problem in discrete time so that the formulation is suitable for working on sampled data. At time 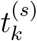, let 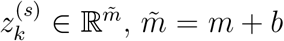, be the concatenation of the external known input *d*^(*s*)^ ∈ ℝ^*b*^ and the observation *y*^(*s*)^ ∈ ℝ^*m*^ sampled at time 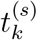 and let 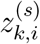 be the *i*^th^ entry of 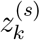. Denote by *N*_*s*_ the number of samples obtained from system *s*, and by *M* the total number of observed systems. Figure 8 illustrates the notation.

**Figure 8:**
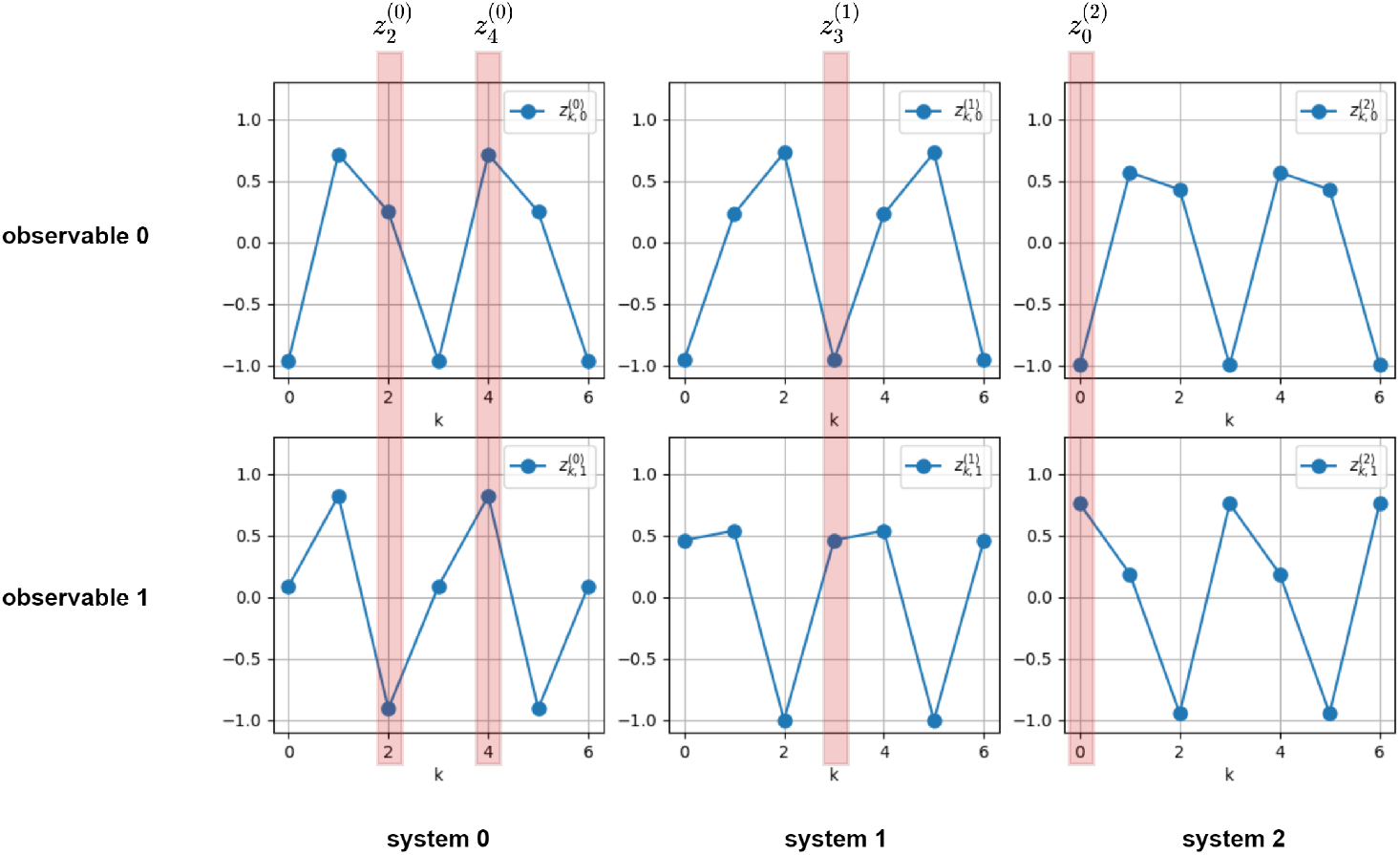
Observed data notations. The k^th^ observation of system *s* is noted by 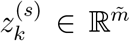 where the i^th^ observable is noted by 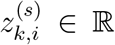. In this illustration the number of observed systems is *M* = 3, the number of samples obtained from each system is *N*_*s*_ = 7 for all systems *s* = 0, …, *M* − 1 and there are two observables, 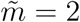.

The methodology we hereby suggest for detecting regulated combinations is based on three key principles. *(i)* A combination that is regulated by the system has a time-dependent structure and is therefore sensitive to shuffles of the time-axis that decouple the components 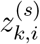 that when coupled form 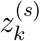 *(ii)* Different observed systems might regulate the same combination, yet about different set-points; *(iii)* A measure for the regulated combination must be universal in the sense that it is independent of physical units.

##### 7.1 Single player optimization

In what follows we construct the optimization algorithm in several methodological stages. *(i)* We begin by formulating a precise mathematical objective that follows the principles of a regulated combination mentioned above, and construct an optimization scheme that minimizes this objective. *(ii)* We then analyze the optimal solutions and show that they fail to capture the regulated combination. *(iii)* To address this shortcoming, in the final stage we construct a two-player optimization scheme that minimizes the same objective yet with respect to a dynamic selection of the observations;

Let 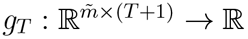 be a function of *T* +1 consecutive observations, *T* ≥ 0, parameterized by *θ*^(*s*)^. For *T* = 0 it is a function of the elements in a single observation. The vector *θ*^(*s*)^ consists of parameters which are specific to system *s*. We refer to *g*_*T*_ (·) as the *regulated combination* and to its output 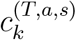 as the *regulated signal* such that for all *k* ≥ *T*,

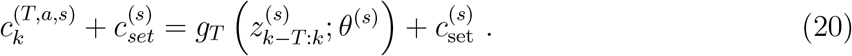

By not including the set-points 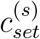 in *g*_*T*_ (·) we follow the second key principle mentioned above, namely, allowing the systems to regulate the same combination, yet about different setpoints for different systems. For the construction of an optimization criterion that derives the *regulated combination* (in the common cases when it is unknown) we should define an objective that obeys the key principles. Intuitively one expects the *regulated signal* to fluctuate about the set-point with ‘small’ deviations, a notion that requires a normalization scheme. A popular normalized measure for the fluctuations of a time-series is the coefficient of variation, CV, which is the standard deviation divided by the mean value. The CV measure does not consider time dependencies in the signal which is our first key principle for a regulated combination. As a result, in many cases one can obtain a combination of two independent observables having an extremely low CV value. In addition, CV is not invariant to the physical units in use as its value is changed by adding an offset (Celsius vs Fahrenheit), a violation of the third principle. We propose to measure the ratio of standard deviations between the *regulated signal* and a signal which is created by evaluating the *regulated combination* on time-shuffled observations.

We begin by examining the output of the *regulated combination* for two types of inputs. The first is the *authentic* input signal that consists of *T* + 1 consecutive observations as in (20). The second input consists of non-consecutive, shuffled observations, that aim to impact the temporal correlations of the original data. We introduce these two sets for each system *s*, corresponding to the two input types. The sets are denoted by Ω^(*T,<*type*>,s*)^, and are constructed as follows.

The first, authentic dataset, consists of the original observed data,

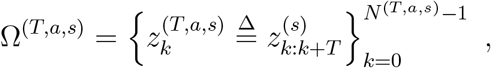

where *N* ^(*T,a,s*)^ = *N*_*s*_ − *T*. The second dataset, referred to as the *‘naively shuffled dataset’*, and denoted by Ω^(*T,n,s*)^, is composed of elements 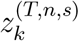, each being a concatenation of nonconsecutive bins. For 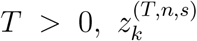 contains a sequence of *T* consecutive bins from the original data, followed by a bin drawn uniformly from the time-series 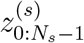. This destroys the correlations between the (*T* + 1)^t*h*^ bin and the preceding *T* bins. For *T* = 0, 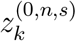 has each and every entry 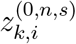 drawn uniformly from 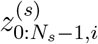. Figure 9 illustrates the composition of these sets. Formally,

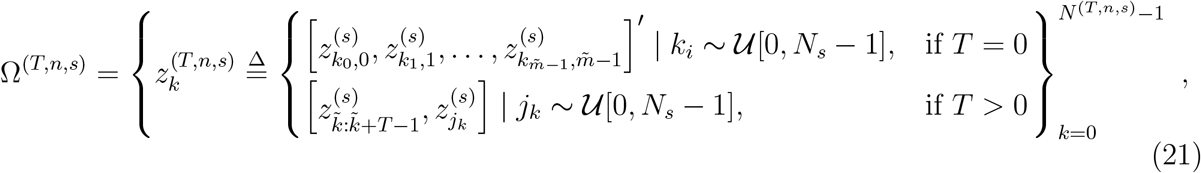

where 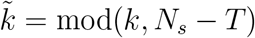 and *N* ^(*T,n,s*)^ ≫ *N* ^(*T,a,s*)^. Note that the dataset Ω^(*T,n,s*)^ is stochastic, and is thus not limited in size to *N* ^(*T,a,s*)^, the number of members in the authentic set. To obtain sufficient variety in the shuffled data we suggest setting *N* ^(*T,n,s*)^ to be an order of magnitude higher than *N* ^(*T,a,s*)^. Denote the distribution of members in the *authentic set* by 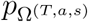, and of members in the *naive shuffle set* by 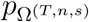, and note that by definition the support of 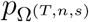 contains the support of 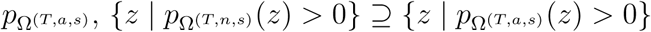.

**Figure 9:**
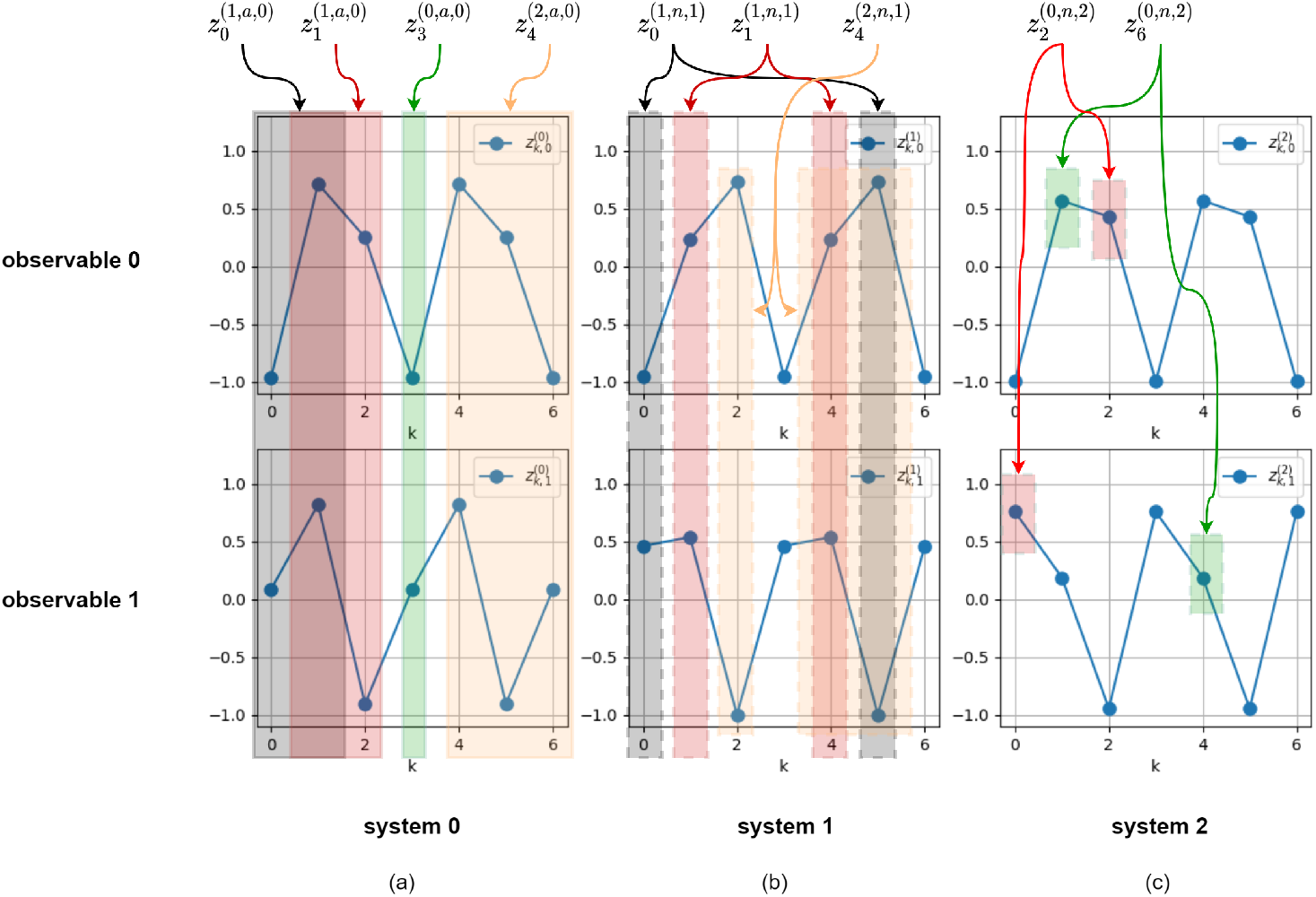
Illustration of the composition of the sets. **(a)** Examples of members 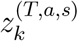 of the *authentic sets* Ω^*T,a,s*^. A member 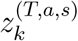 has *T* + 1 consecutive samples starting at sample *k*. **(b)** Example of members 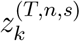 of the *naive shuffle sets* Ω^*T,n,s*^ for *T* > 0. A member 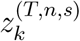 has *T* consecutive samples starting at sample *k* followed by a sample from a random time point. **(c)** Example of members 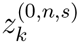 of the *naive shuffle sets* Ω^0,*n,s*^. A member 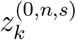 has each observable from a random time point.

We continue by examining the output of *g*_*T*_ (·), the *regulated combination*, for two types of inputs,

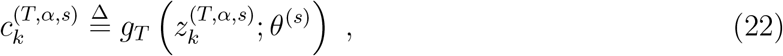

either when evaluated on members of the *authentic set* (*α* = *a*) or when evaluated on members of the *naive shuffle set* (*α* = *n*). Since *g*_*T*_ (·) is a *regulated combination*, and following the principle by which a regulated combination has a time-dependent structure, the time-series 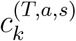 is expected to have a smaller standard deviation than the time-series 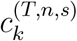, in which the time dependency of the regulated signal was violated by the shuffle. Figure 10 illustrates the time-series at the output of the regulated combination *g*_*T*_ (·) in both cases. We can now form an optimization problem to obtain the combination that is most sensitive to the time shuffle with respect to the ratio of standard deviations. We refer to this optimization as the *‘single player’* because it involves only the *‘combination player’*. As is shown below this optimization turns out not to yield a signal that the system regulates.

**Figure 10:**
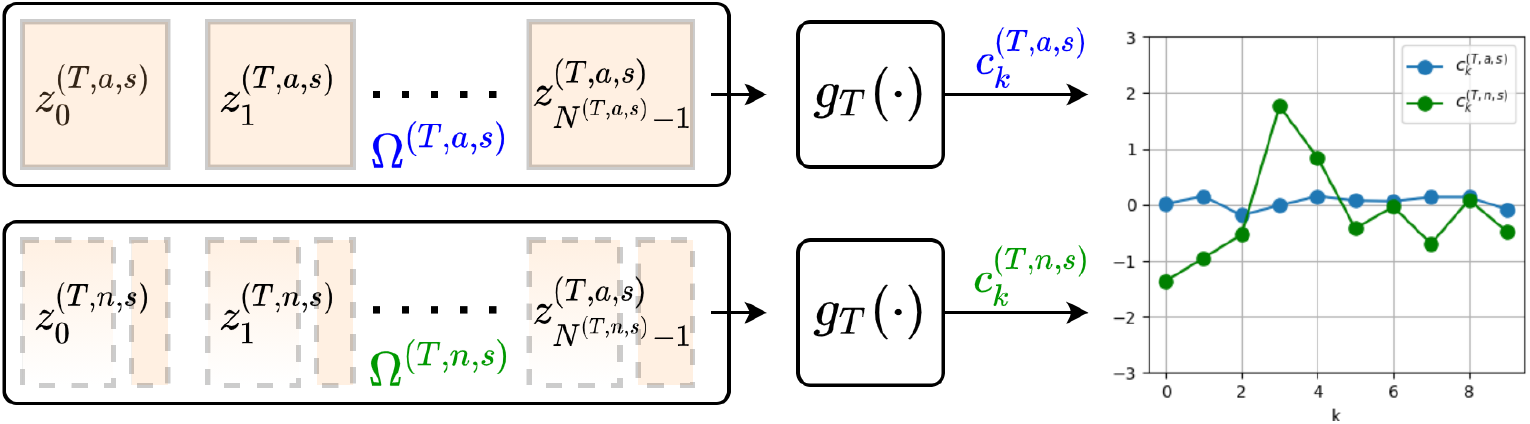
Illustrating the time-series at the output of the regulated combination *g*_*T*_ (·) when evaluated on members of the *authentic set* Ω^(*T,a,s*)^ and on the *naive shuffled set* Ω^(*T,n,s*)^. The shuffled data violates the relations between data-points resulting in an increased standard deviation of the corresponding time-series.

#### Single-player formulation

*Let* 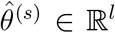, *be a vector of estimated parameters of system s and let* 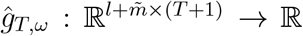 *be a function parameterized by the parameter vector ω. The function* 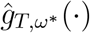 *of the regulated combination is the solution of the optimization problem that is qualitatively written as*,

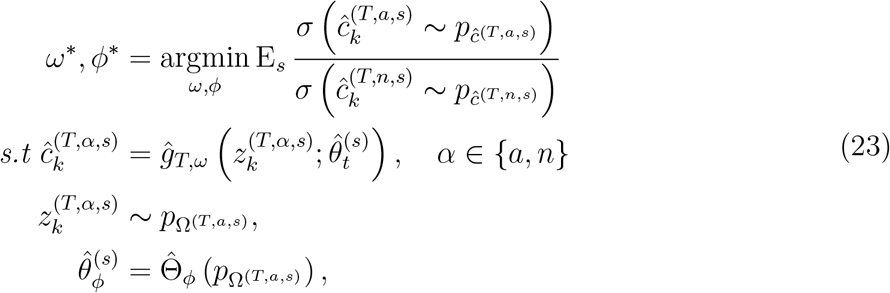

*and exactly, for a given dataset, as*,

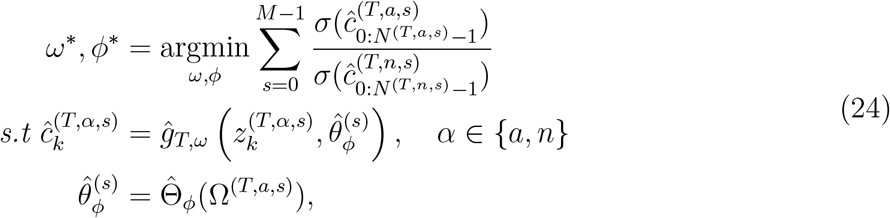

*where* 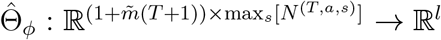 *is a function parameterized by the parameters vector ϕ*.

Optimization problem (24), illustrated in figure 11, has numerous different solutions that yield zero objective value, among which is the solution

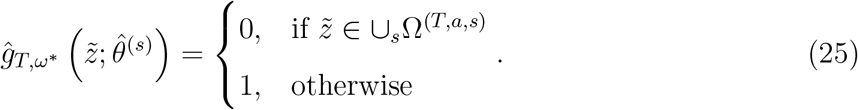

Solution (25) is far from representing a combination that yields a biologically-plausible regulated signal since it is simply 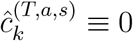. While we are looking for a combination that represents an active feedback mechanism, optimization (24) yields a solution that represents a feature that the authentic time-series always obeys while the time-shuffled series do not.

**Figure 11:**
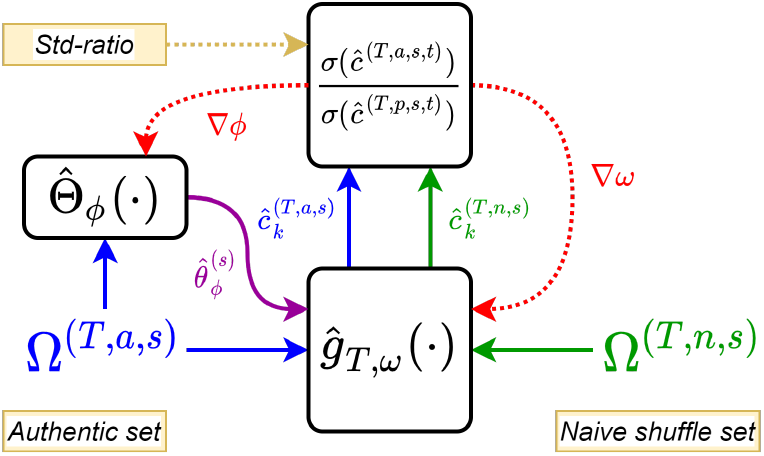
Block diagram of optimization (24). The function *ĝ*_*T,ω*_(·) is optimized to yield the minimal std-ratio, averaged over all systems *s*, between the time-series 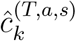 and 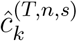 which are its outputs when evaluated on members of the *authentic set* Ω^(*T,a,s*)^ and the *naive shuffle set* Ω^(*T,n,s*)^ respectively. The function 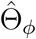, depending on parameters *ϕ*, is used to estimate the parameters 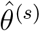 based on the authentic inputs. The gradients, ∇_*ϕ*_ and ∇_*ω*_, contribute to the learning process with respect to the std-ratio objective.

##### 7.2 Two player optimization

In Section 7.1 we translated the principle by which a time-regulated combination has a structure that is sensitive to shuffles of the time axis to a single-objective optimization that seeks to minimize the ratio of the standard deviations, (24). We showed that this optimization scheme does not yield a combination that the system indeed regulates, but, rather, a combination that merely represents a feature that is always obeyed by the data. Analyzing its failure allows us to identify a way to correct it: if we constrain the time-shuffling to include only shuffles that are plausible in light of the data distribution, we may prevent the optimization algorithm from constructing artefact functions that rely solely on structural difference between the measured and shuffled ensembles. In what follows we construct an objective that yields a biologically plausible combination. We do so by constructing IRAS, a two-player algorithm (Algorithm 1) in which one player aims to minimize the ratio of standard deviations, while the second player seeks to keep the optimized combination biologically plausible. To do so we introduce an adaptive shuffle scheme based on the naive shuffle set introduced above.

Recall that in each iteration *t* of the game, the *‘naive shuffle set’*, 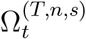, is constructed stochastically by (21) - Algorithm 1 line 8. To construct the new shuffle set, a *‘shuffle player’* examines the *‘naive shuffle set’*, and, according to some fixed policy Π (to be described below), creates a subset, 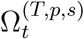,

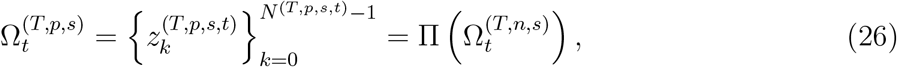

which we refer to as the *‘plausible shuffle set’*, and which has *N* ^(*T,p,s,t*)^ members denoted by 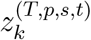 -Algorithm 1 line 9. Once the *‘plausible shuffle set’* set is created, a *‘combination player’* has the objective of minimizing the mean (over all systems) of the ratio of standard deviations, as in (24), except that now the *regulated combination* is evaluated on members of the *‘plausible shuffle set’*, 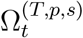, wheres in (24) it was evaluated on members of the *‘naive shuffle set’*, 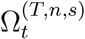. In Algorithm 1, lines 10 − 11, the *combination player* evaluates the current combination 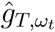 on the two sets, in line 13 it averages the std-ratio over all systems and in line 14 it updates the *regulated combination* to yield 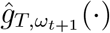 and 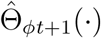.

###### Algorithm 1 IRAS

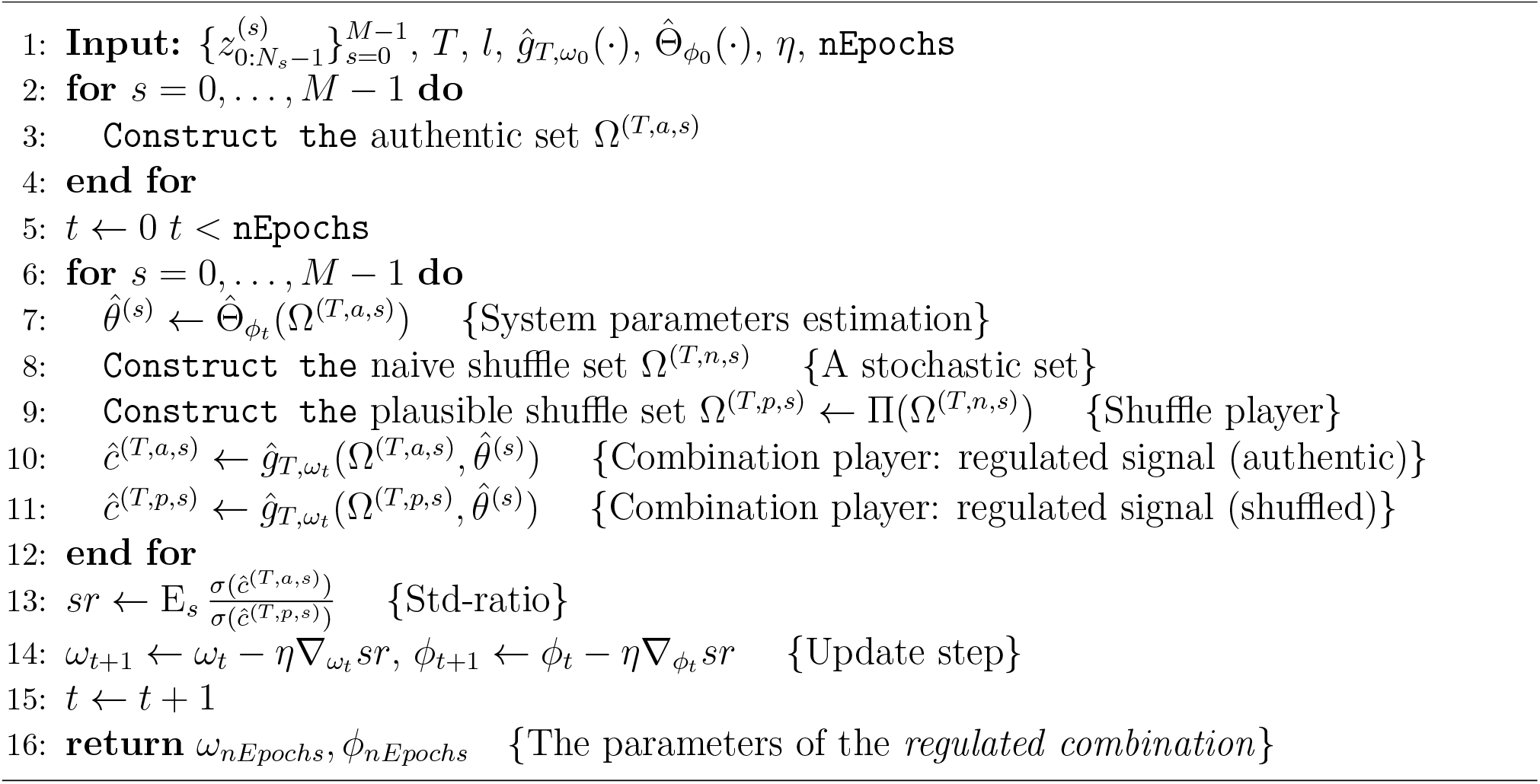

In Section 7.1 we analyzed the case where the policy of the *shuffle player* is to leave the *naive shuffled set* untouched such that 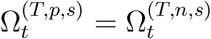, and showed that such a policy yields a combination that represents the feature that is most strongly maintained by the authentic time-series, while being violated by the shuffled time-series. To confront this problem, assume that the *combination player* has learned a combination 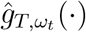 whose violation is biologically non-plausible, thus, a feature that the feedback controller in the system does not operate to maintain. The *combination player* has learned such a feature because the *naive shuffle set* contains members that violate it, that is, members that are not biologically plausible. To derive biologically plausible features, namely features that the controller actively works to regulate, we would like the *shuffle player* to *eliminate* the biologically non-plausible members from the *naive shuffle set*. This encourages the *combination player* to learn the combination that is actively maintained by the controller in the system. To do so we set the shuffle policy Π such that the *plausible shuffle set* resembles the *authentic set* with respect to the combination *ĝ*_*T,ω*_ while maintaining the differences between the naive and authentic sets. Technically, the shuffle player has an **information constraint** because it only has access to values under the combination *ĝ*_*T,ω*_(*z*). Therefore, it outputs data-points distributed by 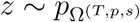 such that

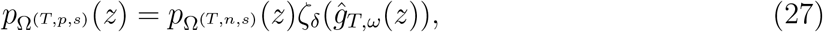

where *ζ*_*δ*_(·) : ℝ → ℝ is a function parameterized by *δ*. Figure 12**a** illustrates the two-player optimization where the objective of the *combination player* is,

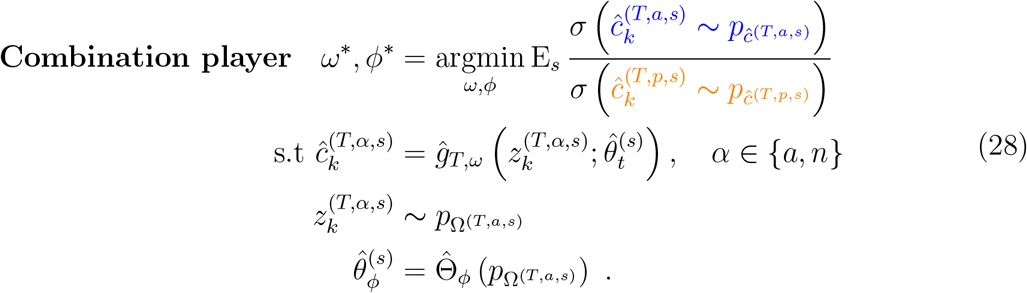

**Figure 12:**
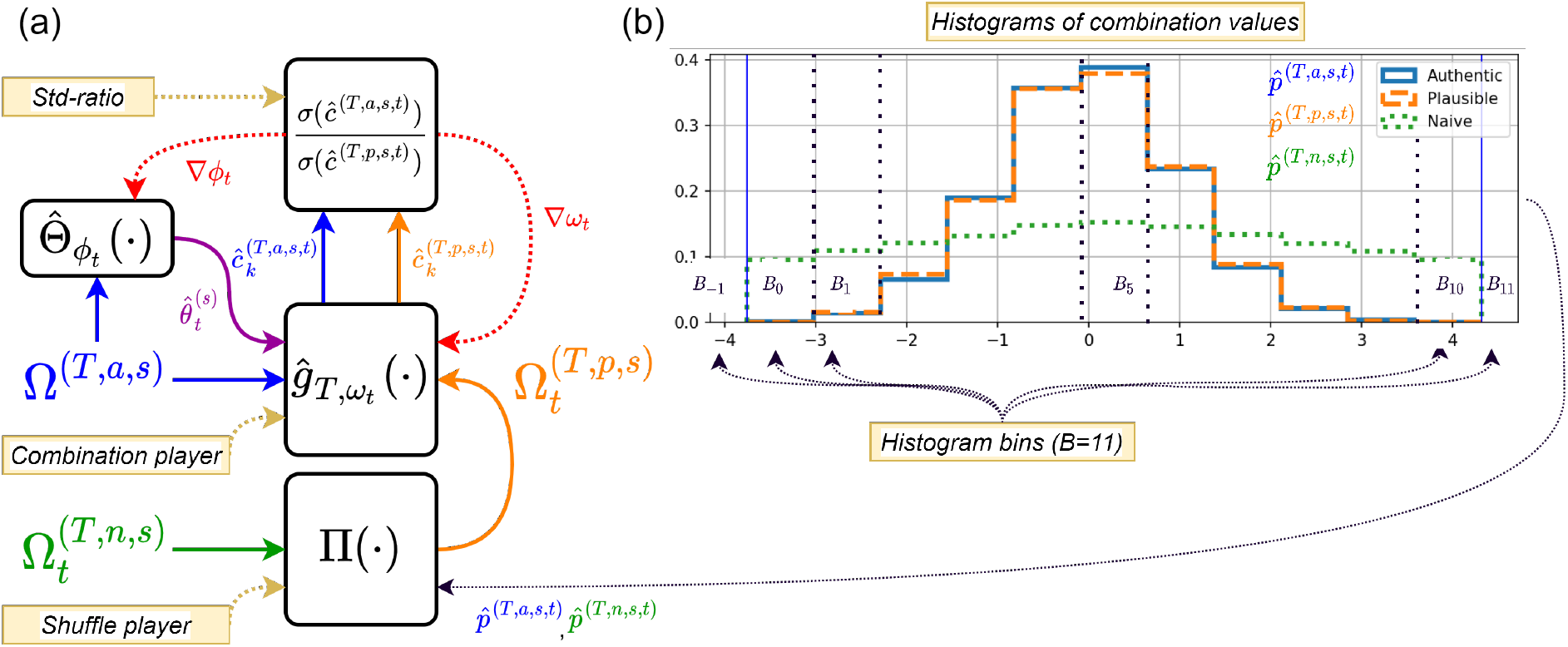
The optimization flow in iteration *t*. **(a)** The *shuffle player* examines the histograms 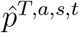 and 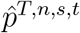 of combination values derived from the *authentic set* and the *naive shuffle set* respectively and constructs the *‘plausible shuffle set’* 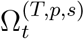. The combination player 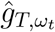 calculates the time-series 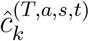 and 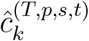 and its weights, *ω*_*t*_, and the weights *ϕ*_*t*_ of the function 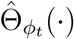 (that estimates the system parameters 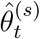) are updated as to minimize the std-ratio. **(b)** Histograms of 13 bins with borders according to (34). The histogram 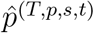of combination values evaluated on the *plausible shuffle set* resembles the histogram 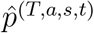 of combination values evaluated on the *authentic set* while the histogram 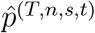 that corresponds to the *naive shuffle set* is significantly wider.

The objective of the *shuffle player* is

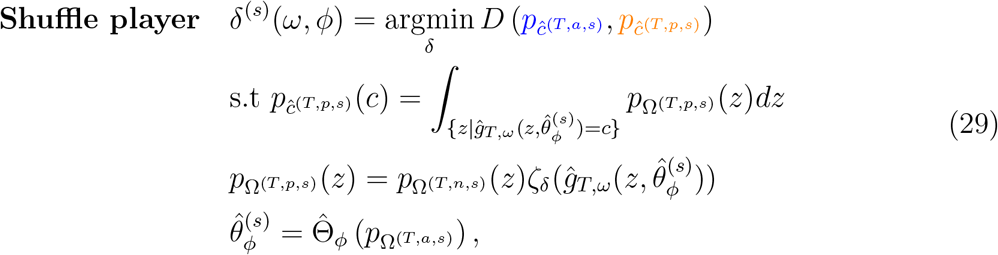

where *D*(·, ·) is a distributional distance metric and *ζ*_*δ*_(·) : ℝ → ℝ is a function parameterized by *δ*, as in (27). To solve the joint optimization problem (28-29) we use the iterative algorithm, Algorithm 1. Let the estimated combination be 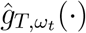, a function parameterized by the vector *ω*_*t*_, where *t* is the optimization-iteration index. For a given policy Π, a gradient-descent update step of the two-players optimization algorithm is given by,

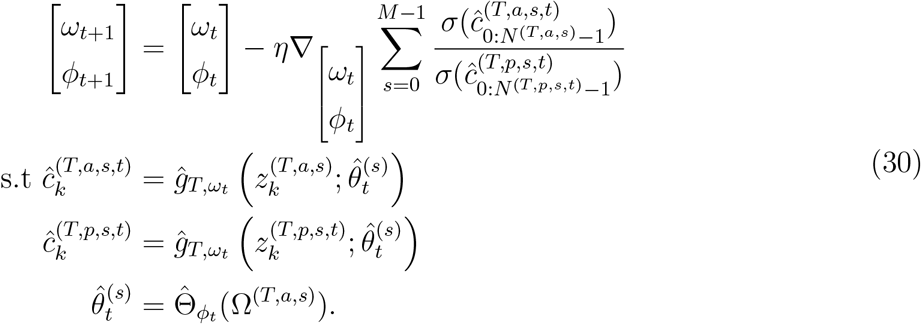

We are left with describing the technical details of the policy Π that creates the *plausible shuffle set* that resembles the *authentic set* with respect to the combination *ĝ*_*T,ω*_.

###### Claim

The optimal solution to the shuffle player’s optimization problem (29) is

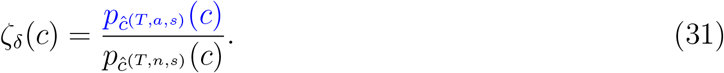

*Proof*.

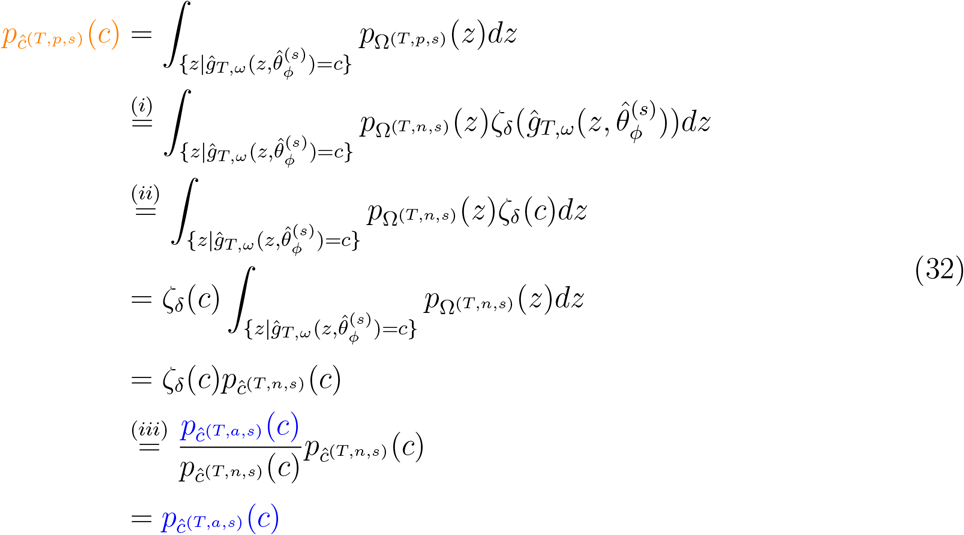

where in (*i*) we substitute (27), in (*ii*) we inserted the projection, 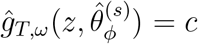 and in (*iii*) we substitute (31). It immediately follows that,

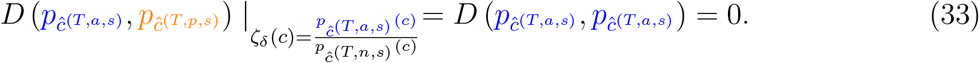

□

According to (29), the policy Π should perform changes to the *naive shuffle set* to obey the constraint by which the distributions of the authentic data and player-shuffled data, with respect to *ĝ*_*T,ω*_, are identical, as manifested in (33). To do so the *shuffle player* estimates these distributions by constructing 1D histograms, and then samples members from the *naive shuffle set* to obtain similar histograms. In each iteration, let the distributions of the combination values of each observed system be approximated by a histogram over the set of bins,

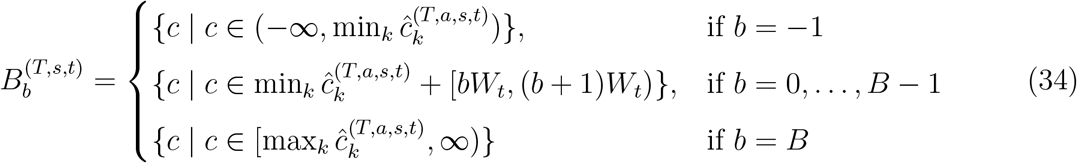

where 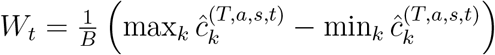 is the width of a single bin in the histogram, and where the number of bins is a user-defined parameter. Then, the histograms of combination values of members in a set are given by,

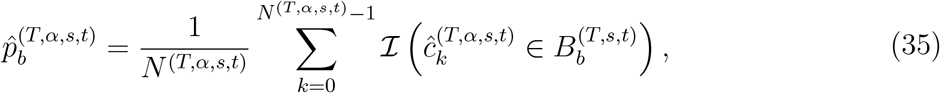

for *α* ∈ {*a, p, s*}, where ℐ(·) is the indicator function. Figure 12**b** depicts the three 1D histograms of the combination 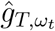, when evaluated on the different sets. In blue, the histogram of authentic data, in green, the naive shuffled data, and in orange, the histogram of player-shuffled data, the outcome of the policy Π manifested in (31).

To block the biologically non-plausible pairs from entering the set of the *‘plausible shuffled pairs’* we designed the policy Π of the *shuffle player* to minimize 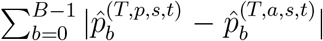, thus creating the *‘plausible shuffle set’* such that the distribution of the combination values of its members resembles the distribution of combination values of members in the *‘authentic set’*. The implementation of (31) is performed by by stochastically blocking a fraction of the members in the *naive shuffle set* from entering the player-shuffled set. This is done by estimating the fraction per each bin in the histogram.

For each combination value *c*, denote by *β*(*c*) the index of the bin in which *c* resides. For each member in the *naive shuffle set*, the *shuffle player* draws a Bernoulli distributed random variable such that,

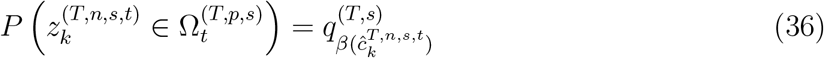

where

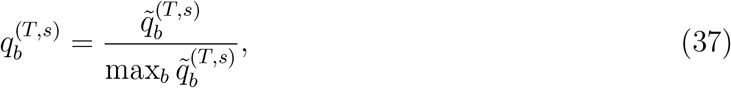

and

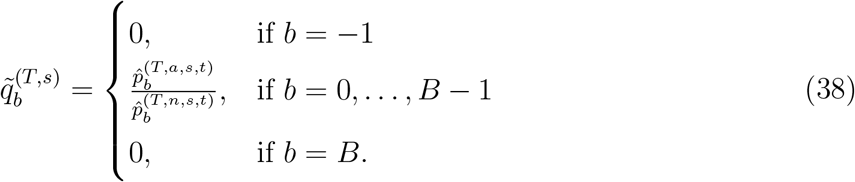

Consider bin *B*5 in figure 12**b**. This bin has the highest ratio between authentic data (blue) and naive shuffled data (green) and thus it serves to normalize the Bernoulli parameters of all bins, (37). All members of the *naive shuffle set* that reside in bin *B*5 are transformed to the player-shuffled set since its corresponding Bernoulli parameter is 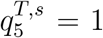. The Bernoulli parameters for all other bins is calculated by evaluating the ratio (38), between the blue and green histograms and normalizing by (37). We note that since Ω^(*T,n,s*)^ ⊃ Ω^(*T,a,s*)^ it is guaranteed that 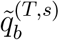 is well defined.

### 8 Implementation details

*ĝ*_*T,ω*_(·). The function *ĝ*_*T,ω*_ implemented as a feed-forward artificial neural network. The input, 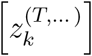or 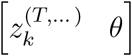 when a parameter vector is estimated as well, is connected to all 32 neurons of the input layer which are themselves connected to a hidden layers having 16 neurons. The output layer has a single neuron which outputs the value of the regulated signal 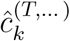 that corresponds the input. The activation function of all neurons is Leaky-ReLU except for the output neuron whose activation function is the Sigmoid function.

Θ_*ϕ*_(·). The function Θ_*ϕ*_ implemented as a feed-forward artificial neural network. The input consists of 100 time samples, *z*_100*i*:100(*i*+1)_ for 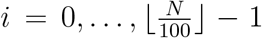, is connected to all 128 neurons of the input layer which are themselves connected to two sequential hidden layers having 128 and 64 neurons each. The output layer has *l* neurons (where *l* is the user-defined number of parameters) which outputs the value *θ*_*i*_. The estimated parameters are given by 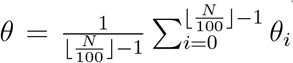. The activation function of all neurons is Leaky-ReLU except for the output neurons which have no activation function.

We simultaneously train the networks as described in Algorithm 1 for a pre-defined fixed number of epochs (500) using the Stochastic-Gradient-descent optimizer with a momentum value of 0.9 and a learning rate of 0.01. See [Zhang et al., 2020] for a detailed explanation of feed-forward neural networks, activation functions and optimizers.

### 9 Algorithm snapshots

Figure 13 is complementary to Figure 4 and displays the combination function and the sampling probability by the shuffle player in different iterations of the algorithm.

**Figure 13:**
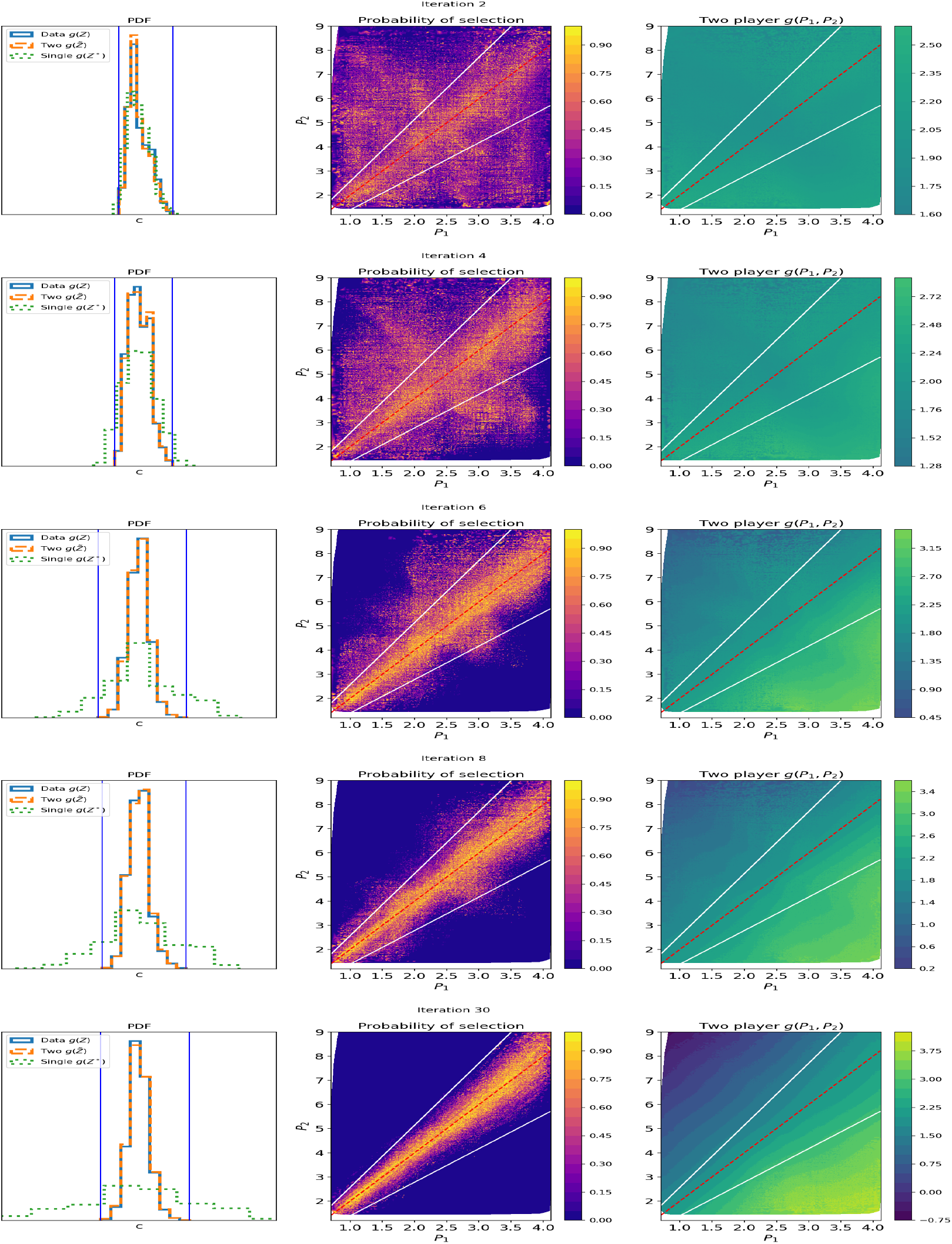
Iterations of the two-player algorithm for the example in Figure 4. The iterations shows a graduate identification of the data region followed by the identification of the control objective, within the region of the data.

### 10 Validation

#### 10.1 Steady states of the kinetic model

At steady state, 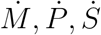 equal zero. Solving the equations gives the steady state

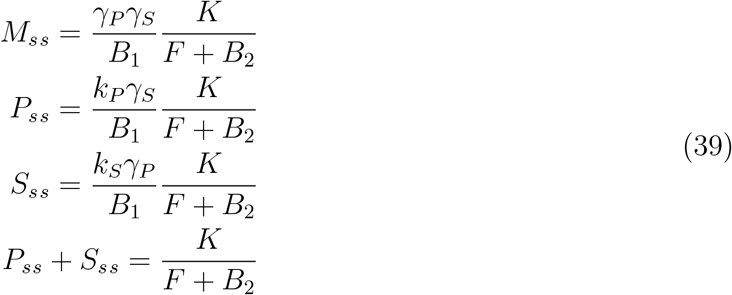

where *B*_*1*_ = *k*_*PγS*_ + *k*_*SγP*_ and 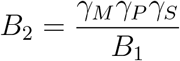. For *F* ≫ *B*_2_, *P*_*ss*_ + *S*_*ss*_ is robust to perturbations in the parameters while the steady states of the three proteins are sensitive to perturbations.

#### 10.2 High and low perturbations in *γ*_*M*_ give rise to The different output combinations

The most conserved combination in the kinetic model depends on the values of the parameters and the perturbations in them. We will consider the cases of low and high perturbations in *γ*_*M*_ relative to the other parameters (*F* and *K* are constant). In the first case, the combination obtained incorporates the effect of *M*. In the second case, we get a behavior that is qualitatively similar to disabling the controller and a combination that reflects the resultant positive correlation between *S* and *P*. The qualitatively different combinations obtained for high and low perturbations in *γ*_*M*_ reflect the two ends of the spectrum of conserved combinations.

##### Low perturbations in *γ*_*M*_

At steady state, and for a single parameter set, 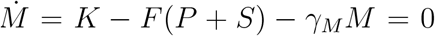. Thus, *F* (*P*_ss_ + *S*_ss_) − *γ*_*M*_ *M*_ss_ = *K*. For small perturbations in *γ*_*M*_, a conserved combination that incorporates *M* is obtained,

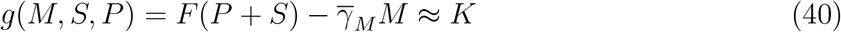

where 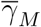 is the mean of *γ*_*M*_ across all realizations. Figure 14A (left panel) shows that this combination is indeed conserved for low perturbations in *γ*_*M*_. Running the algorithm for 30 realization of the model with low perturbations in *γ*_*M*_ gives rise to the combination in (40). The output of the algorithm *g*(*M, S, P*) overlaps with the combination in (40) (Fig. 14A right panel) with a Pearson correlation of 0.97 ± 0.008. Moreover, fitting the output of the network to a linear function of *M, S* and *P* using multivariate linear regression, yields the combination *g*(*M, S, P*) = *S* + *P* + 0.15*M*. Interestingly, the coefficient of *M* in the linear combination is a very close approximation of the ratio 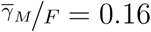.

**Figure 14:**
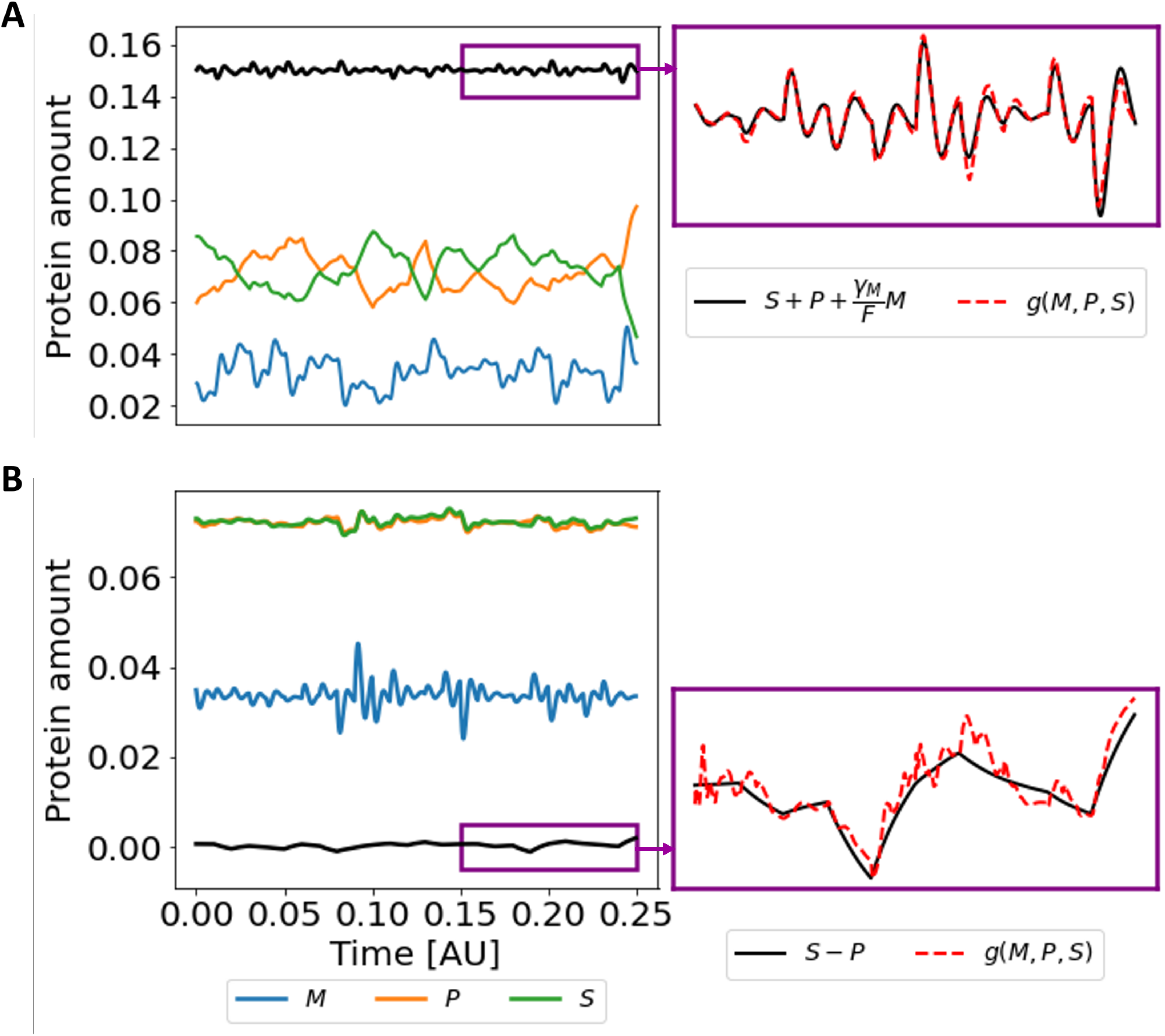
High and low perturbations in *γ*_*M*_ give rise to different output combinations. (A) Low perturbations in *γ*_*M*_ relative to the other parameters. Left Panel: The trajectories of the three proteins and the combination 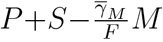 over time. The perturbations in the parameters *γ*_*M*_, *γ*_*S*_, *γ*_*P*_, *k*_*P*_, and *K*_*S*_ were created as described in Figure 5B, where *γ*_*M*_ = 320 ± 5,*γ*_*S*_, *γ*_*P*_ = 70 ± 15,*k*_*S*_, *k*_*P*_ = 150 ± 30. Right Panel: A zoom-in for the combination 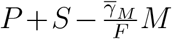 within the purple box along with the output of the algorithm. (B) High perturbations in *γ*_*M*_ relative to the other parameters. Left Panel: The trajectories of the three proteins and the combination *S* − *P* over time, where *γ*_*M*_ = 320 ± 100, *γ*_*S*_, *γ*_*P*_ = 70 ± 1,*k*_*S*_, *k*_*P*_ = 150 ± 1. Right Panel: A zoom-in for the combination *S* − *P* within the purple box along with the output of the algorithm.*F* = 2000, *K* = 300.

##### High perturbations in *γ*_*M*_

High perturbations in *γ*_*M*_, relative to the other parameters, drives the change in M dominantly and in turn affects considerably 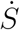 and 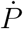. This results in a high positive correlation between S and P since they both follow similar first order kinetics and are both affected by the same source *M* (Figure 14 B left panel). Thus, we expect the output of the algorithm to reflect this positive correlation.Indeed, the output of the algorithm for 3o realization of the model follows the combination *S* − *P* (Figure 14 B right panel).The Pearson correlation between both combinations is 0.79 ± 0.09. Fitting the output of the network to a linear function of *M, S* and *P* using multivariate linear regression, yields the combination *g*(*M, S, P*) = *S* − *P* − 0.06*M*. In fact, this case resembles the case of disabling the controller where M induces the production of S and P without receiving a feedback of their sum.

#### 10.3 Relational dynamics in perception

The estimation errors along a complete trial sequence for various values of *T* are depicted in figure 15. The corresponding mean estimation errors over all trials were depicted in figure **??**.

**Figure 15:**
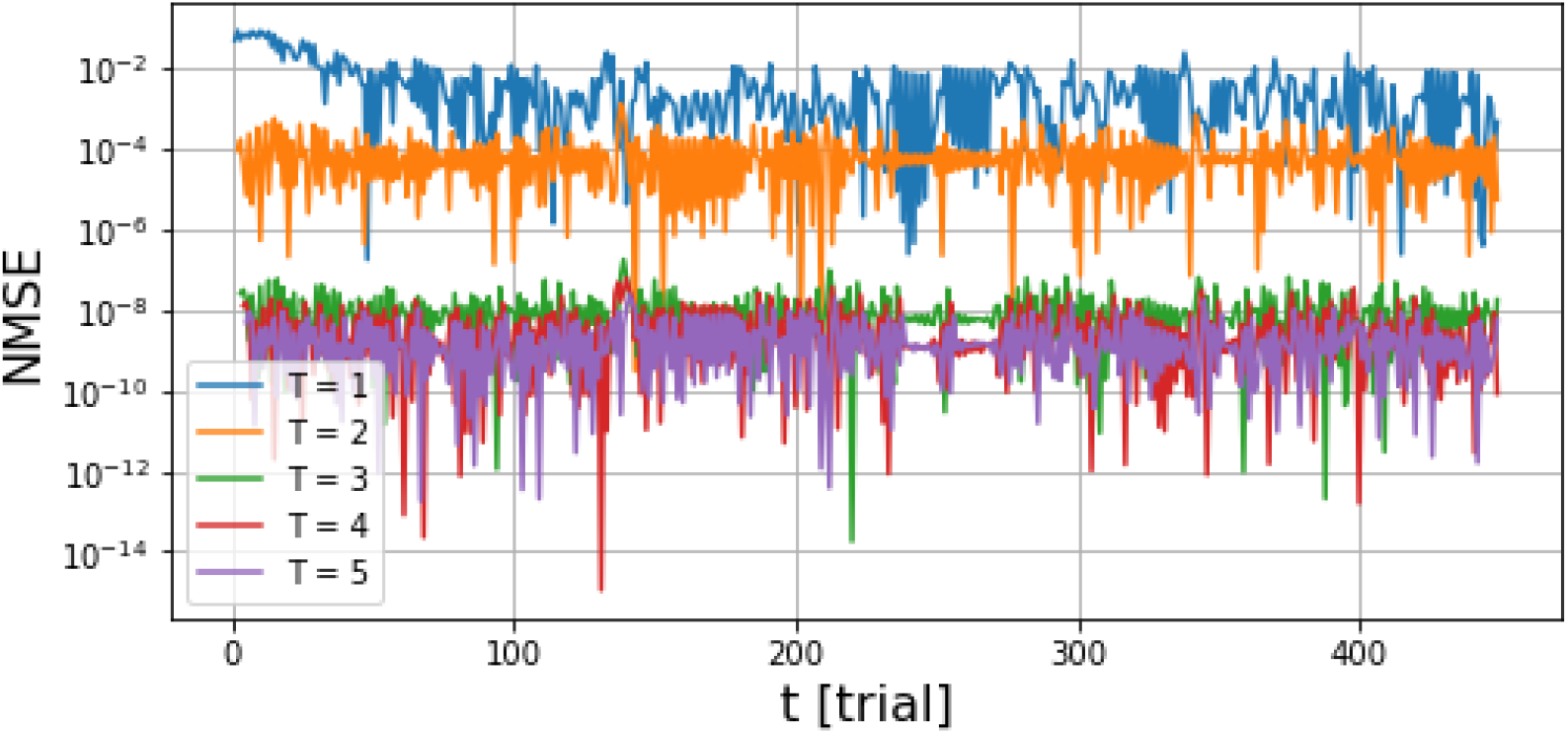
Normalized mean-square-errors of stimuli estimation values along a complete trial sequence. The stimuli estimation is obtained from the analytical expression of the feedback loop the algorithm detected. The estimation errors decrease monotonously up to *T* = 5 implying an effective time-scale of 5 trials.

## Footnotes

* For a detailed view of the iterations see Figure 13 and the video at [iterations.avi](https://www.dropbox.com/s/0hixr4bzohswcwd/shuffleSelectionProb.avi?dl=0)

* Observability, in the sense of Control Theory, refers to the ability of determining the internal state of a system given measurements, e.g. Chapter 1.7. in [Simon, 2006]

## References

Alet, F., Doblar, D., Zhou, A., Tenenbaum, J., Kawaguchi, K., and Finn, C. (2021). Noether networks: meta-learning useful conserved quantities. Advances in Neural Information Processing Systems, 34:16384–16397.

Åström, K. J. (2012). Introduction to stochastic control theory. Courier Corporation.

Åström, K. J. and Murray, R. M. (2021). Feedback systems: an introduction for scientists and engineers. Princeton university press.

Billman, G. E. (2020). Homeostasis: the underappreciated and far too often ignored central organizing principle of physiology. Frontiers in Physiology, page 200.

Chang, M. B., Ullman, T., Torralba, A., and Tenenbaum, J. B. (2016). A compositional object-based approach to learning physical dynamics. arXiv preprint arXiv:1612.00341.

Chen, B., Huang, K., Raghupathi, S., Chandratreya, I., Du, Q., and Lipson, H. (2022). Automated discovery of fundamental variables hidden in experimental data. Nature Computational Science, 2(7):433–442.

Cosentino, C. and Bates, D. (2011). Feedback control in systems biology. Crc Press.

Daniels, B. C. and Nemenman, I. (2015). Automated adaptive inference of phenomenological dynamical models. Nature communications, 6(1):1–8.

de Avila Belbute-Peres, F., Smith, K., Allen, K., Tenenbaum, J., and Kolter, J. Z. (2018). End-to-end differentiable physics for learning and control. Advances in neural information processing systems, 31.

El-Samad, H. (2021). Biological feedback control—respect the loops. Cell Systems, 12(6):477–487.

Faisal, A. A., Selen, L. P., and Wolpert, D. M. (2008). Noise in the nervous system. Nature reviews neuroscience, 9(4):292–303.

Goodfellow, I., Pouget-Abadie, J., Mirza, M., Xu, B., Warde-Farley, D., Ozair, S., Courville, A., and Bengio, Y. (2014). Generative adversarial nets. Advances in neural information processing systems, 27.

Greydanus, S., Dzamba, M., and Yosinski, J. (2019). Hamiltonian neural networks. Advances in neural information processing systems, 32.

Haber, A. and Schneidman, E. (2022). Learning the architectural features that predict functional similarity of neural networks. Physical Review X, 12(2):021051.

Hamrick, J. B., Allen, K. R., Bapst, V., Zhu, T., McKee, K. R., Tenenbaum, J. B., and Battaglia, P. W. (2018). Relational inductive bias for physical construction in humans and machines. arXiv preprint arXiv:1806.01203.

Hsiao, V., Swaminathan, A., and Murray, R. M. (2018). Control theory for synthetic biology: recent advances in system characterization, control design, and controller implementation for synthetic biology. IEEE Control Systems Magazine, 38(3):32–62.

Kotas, M. E. and Medzhitov, R. (2015). Homeostasis, inflammation, and disease susceptibility. Cell, 160(5):816–827.

Lancaster, G., Iatsenko, D., Pidde, A., Ticcinelli, V., and Stefanovska, A. (2018). Surrogate data for hypothesis testing of physical systems. Physics Reports, 748:1–60.

Liu, Y.-Y., Slotine, J.-J., and Barabási, A.-L. (2013). Observability of complex systems. Proceedings of the National Academy of Sciences, 110(7):2460–2465.

Ljung, L. (1998). System identification. In Signal analysis and prediction, pages 163–173. Springer.

Marom, S. (2010). Neural timescales or lack thereof. Progress in neurobiology, 90(1):16–28.

Marom, S. and Wallach, A. (2011). Relational dynamics in perception: impacts on trial-to-trial variation. Frontiers in Computational Neuroscience, 5:16.

Monto, S., Palva, S., Voipio, J., and Palva, J. M. (2008). Very slow eeg fluctuations predict the dynamics of stimulus detection and oscillation amplitudes in humans. Journal of Neuroscience, 28(33):8268–8272.

Reichl, L. E. (1999). A modern course in statistical physics.

Sakurai, J. J. and Commins, E. D. (1995). Modern quantum mechanics, revised edition.

Santoro, A., Raposo, D., Barrett, D. G., Malinowski, M., Pascanu, R., Battaglia, P., and Lillicrap, T. (2017). A simple neural network module for relational reasoning. Advances in neural information processing systems, 30.

Schmidt, M. and Lipson, H. (2009). Distilling free-form natural laws from experimental data. science, 324(5923):81–85.

Shen, J., Liu, F., Tu, Y., and Tang, C. (2021). Finding gene network topologies for given biological function with recurrent neural network. Nature communications, 12(1):1–10.

Simon, D. (2006). Optimal state estimation: Kalman, H infinity, and nonlinear approaches. John Wiley & Sons.

Stawsky, A., Vashistha, H., Salman, H., and Brenner, N. (2021). Multiple timescales in bacterial growth homeostasis. bioRxiv.

Tegnér, J. and Björkegren, J. (2007). Perturbations to uncover gene networks. TRENDS in Genetics, 23(1):34–41.

Tenenbaum, J. B., Silva, V. d., and Langford, J. C. (2000). A global geometric framework for nonlinear dimensionality reduction. science, 290(5500):2319–2323.

Watters, N., Zoran, D., Weber, T., Battaglia, P., Pascanu, R., and Tacchetti, A. (2017). Visual interaction networks: Learning a physics simulator from video. Advances in neural information processing systems, 30.

Zhang, A., Lipton, Z. C., Li, M., and Smola, A. J. (2020). Dive into Deep Learning. https://d2l.ai.

